# *BpLAZY1A* Mediates Transcriptional Polarity and Drives Adaxial Tension Wood–Like Tissue Formation in Silver Birch (*Betula pendula*) Branches

**DOI:** 10.64898/2026.09.12.750306

**Authors:** Sampo Muranen, Àngela Carrió-Seguí, Ema Marmara, Aleš Pěnčík, Juha Immanen, Ilhan Duru, Mira Viljanen, Heikki Suhonen, Kirsi Svedström, Xueping Shi, Maja Ilievska, Pasi Rastas, Chang Su, Hanna Koivula, Juan Alonso-Serra, Ondřej Novák, Pawel Roszak, Kaisa Nieminen, Ykä Helariutta

## Abstract

A defining feature of most trees is the architectural distinction between a vertically growing main stem and laterally growing branches. This growth habit increases fitness by increasing photosynthetically active surface area and shading competitors. Here, we studied the weeping birch cultivar *Betula pendula* ‘Youngii’ to uncover the mechanisms that maintain lateral branch growth. We identified a loss-of-function mutation in *BpLAZY1A*, a core component of the gravitropic signalling pathway, as the cause of the weeping phenotype.

Forward genetic analysis demonstrated that the weeping phenotype is recessive in silver birch. Transgenic *BpLAZY1A* lines phenocopied ‘Youngii’, confirming the functional role of *BpLAZY1A*. Reporter analysis revealed that *BpLAZY1A* is expressed predominantly in gravity-sensing starch sheath cells, with occasional expression in the main stem phloem. Time-lapse imaging revealed two distinct gravitropic responses during branch development -an early response associated with establishment of the branch apex gravitropic set-point angle, and a later response associated with polar reinforcement at the branch base. The main stem retained normal gravitropic responses in *BpLAZY1A RNAi* lines, indicating a branch-specific role for *BpLAZY1A*.

Integrative analyses combining transcriptomics, chemical profiling, and histochemistry indicated that *BpLAZY1A* establishes adaxial–abaxial polarity during early branch development. In wild-type branches, pectin-rich tension wood-like tissue formed preferentially in the adaxial xylem, accompanied by adaxial expression of pectin- and tension wood-associated genes, including *BpRRT1* and *BpCOBRA-LIKE4*. Auxin-related genes, including *BpIAA29* and *BpSAURs*, were preferentially upregulated in the abaxial side. This transcriptional asymmetry was reduced in *BpLAZY1A RNAi* line 1, where tension wood-like tissue formed in both adaxial and abaxial sides of the xylem, indicating a loss of polarity.

Together, these findings demonstrate that *BpLAZY1A* functions specifically in branch gravitropism and links spatially asymmetric gene expression with branch growth orientation and biomechanical reinforcement. Our results identify *BpLAZY1A* as a key regulator coordinating gravitropic signalling, tissue polarity, and the developmental biomechanics underlying lateral branch growth.

## Introduction

Gravitropism is an adaptive mechanism that determines plant architecture by directing organ growth relative to gravity. In vascular plants, shoots generally exhibit negative gravitropism by growing upward to maximize light capture, whereas roots display positive gravitropism, providing anchorage and efficient acquisition of water and nutrients (Morita, 2010). This coordinated growth has been crucial for the evolutionary success of land plants and their adaptation to diverse environments (Zhang *et al*., 2019).

The first mechanistic explanation of gravity sensing was proposed by Haberlandt (1900), who suggested that starch-filled amyloplasts (statoliths) sediment within specialized gravity sensing cells called statocytes. The Cholodny-Went model, which was devised in the 1920s, proposed that gravitropic and phototropic curvature results from asymmetric auxin redistribution. Differential auxin accumulation produces unequal cell elongation, causing shoots to grow towards light (phototropism) and roots to grow towards gravity (gravitropism) (Went, 1974). Current models describe gravitropism as a four-stage process involving gravity perception, signal transduction, asymmetric auxin transport, and differential growth (Morita *et al*., 2026).

Although this framework explains primary root growth, it does not fully address shoot gravitropism, especially in woody species undergoing extensive secondary growth. In woody branches, gravity-induced auxin gradients likely influence vascular cambium activity and secondary xylem development. Consistent with this idea, inclined poplar stems display elevated auxin signalling in the adaxial xylem and abaxial cortex, indicating tissue-specific auxin distribution (Gerttula *et al*., 2015). These gradients likely regulate secondary growth, cell expansion, and the formation of specialized tissues such as tension wood in a cell type specific manner, also in the branch context.

The characteristic horizontal (plagiotropic) growth orientation of primary branches requires a careful balancing. Branches must resist gravitational forces while preventing a transition to upright (orthotropic) growth. The mechanisms maintaining branch orientation remain poorly understood (Chen *et al*., 2025). In angiosperm trees, displaced main stems produce tension wood that generates forces restoring vertical growth (Wilson and Gartner, 1996). In contrast, tension wood formation is suppressed in two-year-old branches of ash (*Fraxinus americana*) and cherry (Prunus serotina) and becomes prominent after disruption of apical dominance by girdling (Wilson and Archer, 1983; Wilson, 2000). Also, Kohler *et al*. (2024), did not observe tension wood in young branches of peach (*Prunus persica*). These data suggest a distinct regulatory mechanism governing young branch posture.

In *Arabidopsis*, LAZY proteins localize to both amyloplast and plasma membranes of columella cells, linking gravity perception to signalling. Following gravistimulation, MAPK-mediated phosphorylation enhances LAZY association with TOC proteins, allowing movement with sedimenting amyloplasts. At the lower plasma membrane, LAZY recruits RLD proteins, forming a complex that polarizes D6PK and promotes asymmetric PIN redistribution and activation, generating directional auxin flow and root curvature (Furutani *et al*., 2020; Chen *et al*., 2023; Kulich *et al*., 2023; Morita *et al*., 2026). Whether the same pathway regulates shoot gravitropism remains unknown.

Understanding the LAZY-mediated regulation of shoot gravitropism has practical importance because branch angle and shoot architecture strongly influence light interception, canopy structure, lodging resistance, and crop productivity (Hollender and Dardick, 2014). Woody species provide an opportunity to investigate how gravitropic signalling integrates with secondary growth and wood development. Silver birch (*Betula pendula*) is particularly suitable for such studies owing to its short juvenile phase, well characterized diploid genome and substantial architectural diversity (Longman and Wareing, 1959; Salojärvi *et al*., 2017; Su et al., 2023; Fig. S1).

Here, we demonstrate in silver birch that *BpLAZY1A* is a key component of branch gravitropism response, which involves formation of pectin-rich tension wood-like tissue in the adaxial xylem. RNA-seq analyses of branches revealed polarized expression of pectin- and tension wood-related genes on the adaxial side, while auxin-related genes were enriched abaxially. This spatial asymmetry was abolished in *BpLAZY1A RNAi* plants, providing new insight into the genetic regulation of branch posture in woody species undergoing secondary growth.

## Results

### *Betula pendula* ‘Youngii’ is a naturally occurring recessive *Bplazy1a* mutant

Our forward genetics program in *Betula* species has focused on identifying putative monogenic mutants exhibiting distinct shoot architecture phenotypes. Through an extensive search-and-cross program, we have identified several candidates that will provide insight into the genetic regulation of tree shoot system architecture (Fig. S1). Within this program, we have previously identified a naturally occurring, putatively deleterious *Bplazy1* mutation in the birch cultivar *Betula pendula* ‘Youngii’ (Fig. 1A; Salojärvi *et al*., 2017). The *LAZY* gene family comprises six members in Arabidopsis (Yoshihara and Spalding, 2017) and four in silver birch. Notably, silver birch contains two closely related *BpLAZY1* homologs, here designated as *BpLAZY1A* (Bpev01.c0052.g0076.m0001) and *BpLAZY1B* (Bpev01.c0566.g0022.m0001) (Fig. S3A). The causal mutation was identified using whole genome sequencing (WGS) data by screening for disruptive variants in candidate genes involved in the gravitropic pathway. The primary candidate was the closest birch ortholog of *AtLAZY1*, as mutations in this gene and it’s orthologs are associated with *lazy*-like phenotypes in rice, maize, *Arabidopsis*, and apple tree (Li *et al*., 2007; Dong *et al*., 2013; Yoshihara *et al*., 2013; Taniguchi *et al*., 2017; Dougherty *et al*., 2023). This approach highlighted a homozygous single nucleotide polymorphism (SNP) in the third exon of *BpLAZY1A* that introduces a premature stop codon in *B. pendula* ‘Youngii’ (Fig. 1D; Salojärvi *et al*., 2017). To determine the inheritance pattern, *B. pendula* ‘Youngii’ was crossed with wild-type pollen (E1970 from the LUKE collection). All F₁ progeny exhibited a wild-type phenotype (Fig. 1B), consistent with recessive inheritance. A subsequent backcross to the ‘Youngii’ yielded a 1:1 segregation ratio (χ² = 1.11, p = 0.29), confirming that the weeping phenotype of *B. pendula* ‘Youngii’ is controlled by a single recessive locus (Fig. 1C). The association of the point mutation with the weeping phenotype in the segregating population was confirmed with restriction enzyme digestion (Fig. S2).

**Figure 1.**
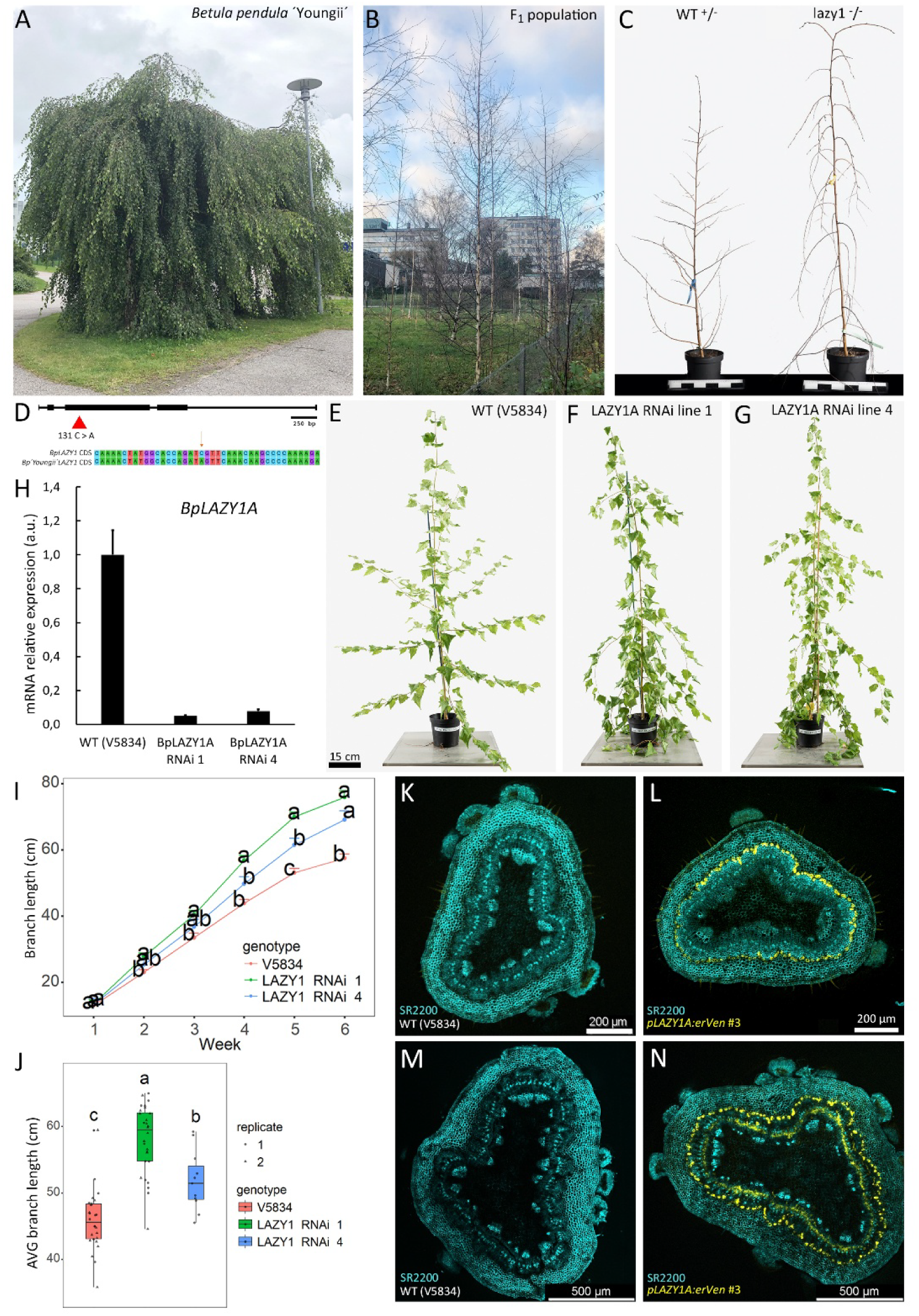
Characterization of the *lazyl* mutation in *Betuia pendula* ‘Youngii’. (A) *B. pendula* ‘Youngii’ is a natural *bpíazyla* mutant. (B) F, progeny from the ‘Youngii’ × E1970 cross display a 100% wild-type phenotype. (C) The BCi population of ‘Youngii’ × Fi consists of 90 individuals and segregates into 50 wild-type and 40 mutant trees, which fits the expected 1:1 ratio (x^2^ = 1.11, p = 0.29), consistent with the inheritance pattern of a single locus recessive mutation. Scale bars 30 cm. (D) *B. pendula* ‘Youngii’ carries a premature stop codon in the third exon of the *BpLAZYlA* gene. (E-G) *BpLAZYlA RNAi* lines phenocopy the weeping phenotype of *B. pendula* ‘Youngii’. (H) *BpLAZYlA* expression is downregulated in the RNAi lines. Data are represented as mean ±1 SD. N = 3 biological replicates. (I) Branches of the *BpLAZYlA RNAi* lines grow faster than the WT {V5834). The difference is statistically significant from week 5 onwards. Different letters indicate significant differences among groups as determined by one-way ANOVA followed by Tuke/s HSD post-hoc test (p < 0.05). Data are represented as mean ±1 SE. N = 8 biological replicates. Three branches {technical replicates) at the base of the stem which were measured once a week per biological replicate. (J) Branches of the *BpLAZYlA RNAi* lines grow longer than the WT (V5834). Different letters indicate significant differences among groups as determined by one-way ANOVA followed by Tukey’s HSD post-hoc test (p < 0.05). All branches were measured per biological replicate. (K) WT (V5834) as an autofluorescence branch control. (L) *BpLAZYlA* expression localizes to the starch sheath (endodermis) of the branch in the 3rd basal internode. (M) WT (V5834) as an autofluorescence main stem control. (N) *BpLAZYlA* expression localizes to the starch sheath and to the inner side of phloem fibers in the main stem.

We conducted a comparative analysis of previously published/available expression data across multiple species exhibiting pronounced *lazy*-like shoot phenotypes (Li *et al*., 2007; Dong *et al*., 2013; Yoshihara *et al*., 2013). This analysis indicated that mutations within the LAZY1/6 clade are consistently associated with the *lazy* phenotype in the shoots (Fig. S3A). Single-cell RNA-seq data of *Arabidopsis* primary root tip support this distinction. *AtLAZY1* is strongly expressed in the endodermis, indicating a potential link to shoot gravitropism, while *AtLAZY2-4* are strongly expressed in the columella cells (root gravitropism). *AtLAZY5* has peak expression in the cortex, while no data was available of *AtLAZY6* expression (Fig. S3B, data from Su *et al*., 2023).

To validate the role of *BpLAZY1A* (mutated in *B. pendula* ‘Youngii’) in branch gravitropism, we generated *BpLAZY1A RNAi* lines using a wild-type laboratory clone (V5834 from the LUKE collection) as the background. These RNAi lines phenocopied the weeping branch phenotype, providing strong evidence that *BpLAZY1A* is required for gravitropic responses in silver birch branches (Fig. 1E–G). RT–qPCR confirmed that *BpLAZY1A* expression is downregulated in the RNAi lines (Fig. 1H).

*BpLAZY1A RNAi* plants displayed accelerated branch apical growth compared to the wild type. The difference in branch length became statistically significant from week 5 onwards (Fig. 1I). *BpLAZY1A RNAi* plants also tended to produce longer branches when measured three months after potting (Fig. 1J). *BpLAZY1A RNAi* plants were also slightly taller, but branch and stem diameter, branch number, and internode number did not display statistically significant difference between wild-type and RNAi lines when measured three months after potting (Fig. S4).

### *BpLAZY1A* is expressed in the starch sheath cells throughout the shoot system

*pBpLAZY1A::erVEN* reporter lines were generated to study the expression pattern of *BpLAZY1A* throughout the shoot system. Tissue sections were obtained from three positions of two-month-old plants that were ∼150 cm tall. The third apical internode, the third basal internode, and the first basal internode were sampled from the branches. The main stem samples were taken from the 3^rd^ apical internode, middle of the plant and 20 cm above the soil. Confocal analysis of these sections demonstrated that *BpLAZY1A* is expressed mainly in the gravity sensing starch sheath (endodermis) throughout the shoot system (Fig. 1K-N, Fig. S5-6). Additionally, we observed *BpLAZY1A* expression also in the phloem in the main stem (Fig. 1N; Fig. S6B-C).

### High-resolution time-lapse imaging reveals spatial and organ specific functions of *BpLAZY1A*

To investigate the spatial and organ-specific roles of *BpLAZY1A*, we performed high-resolution time-lapse imaging at 1-hour imaging intervals under two lighting settings: An 18/6-hour light-dark cycle for the segregating back-crossed/BC1 population, and continuous light for V5834 and *BpLAZY1A RNAi* plants. We carried out the initial time-lapse experiments with the segregating population.

These studies displayed that the 6-hour dark period introduced abrupt apical movements. This was due to the black “nighttime” frames which were removed from the dataset before conversion to videos. To eliminate this artifact, imaging was conducted under continuous illumination for the V5834 vs *BpLAZY1A RNAi* videos. Experiments were performed under three conditions: Straight, tilted, and decapitated growth setups.

Under straight growth conditions (Movie 1), wild-type branch apices exhibited active and regular orthogonal movements during early development (71–90 h). Comparable early orthogonal movement was also observed in the *BpLAZY1A RNAi* line 4 (Fig. 2A–B). At the later stage (231–250 h), orthogonal activity persisted in wild-type branches but was largely absent in both *BpLAZY1A RNAi* lines (Fig. 2C–D). Notably, the basal internode exhibited early bending in both *BpLAZY1A RNAi* and *Bplazy1a* mutant plants, whereas wild-type branch bases remained relatively straight (Fig. 2E–F).

**Figure 2.**
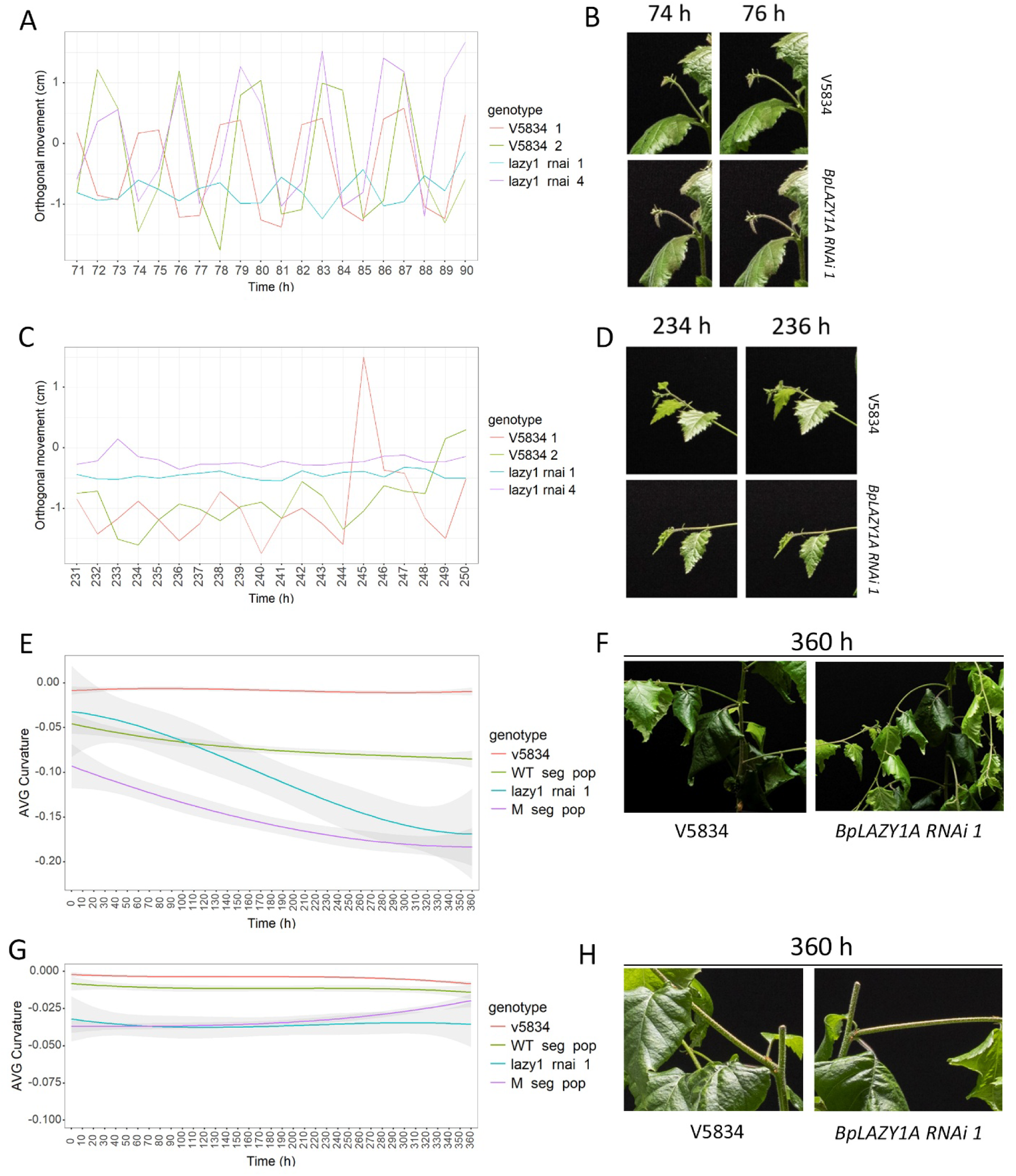
Time-lapse photography characterisation of the *bplazy1a* phenotype. (A) Time-lapse imaging reveals a highly rhythmic back-and-forth movement in the WT and the *BpLAZYl RNAi* line 4 with a periodicity of approximately four hours. This oscillatory pattern is absent in the *BpLAZYlA RNAi* line. **N** = 1 biological replicate per experiment. (B) Close-up images at 74h and 76h highlight the characteristic movement in the WT. (C) Orthogonal movement is apparent at later stages in the WTs but it is almost completely abolished in the *BpLAZYlA RNAi* lines. N = 1 biological replicate per experiment. (D) Close-up images at 234h and 236h highlight the characteristic movement in the WT. (E) Under normal growth conditions, WT branches maintain a low curvature in the basal internode, whereas *BpLAZYlA RNAi 1* and *bplazyla* mutant exhibit pronounced bending. **N** = 3 technical replicates from 1 biological sample. Data are represented as mean ±1 SE. {F) Close-up images higlight the bending of the basal internode in the *BpLAZYlA RNAi* line 1. (G-H) Following decapitation, the basal internodes of the uppermost branches remain straight in WT, *bplazyla* mutant, and *BpLAZYlA RNAi* line 1 demonstrating that decapitation rescues the basal internode phenotype in *BpLAZYlA RNAi* and *bplazyla* mutant. N = 2 technical replicates from 1 biological sample. Data are represented as mean ±1 SE.

Under tilted conditions (Movie 2), both wild-type and *bplazy1a* mutant main stems of the segregating population displayed comparable gravitropic responses. A rapid response (three hours after tilting) was observed in the apical regions, followed by a slower phase, likely reflecting tension wood formation in the main stem. Similar patterns were observed when comparing V5834 and *BpLAZY1A RNAi* plants.

To assess tension wood formation in the absence of *BpLAZY1A*, we examined the effect of decapitation, a condition known to induce tension wood formation in the uppermost branches under the decapitation site (Wilson and Archer, 1983; Wilson, 2000). Time-lapse imaging revealed upward reorientation of the uppermost branches in wild types, *Bplazy1a* mutant and in the *BpLAZY1A RNAi* (Movie 3). This response is likely driven by abrupt tension wood formation, as both V5834 and *BpLAZY1A RNAi* plants displayed extensive tension wood development four weeks after decapitation (Fig. S7A–B). Notably, the apical regions of the mutant branches maintained a weeping phenotype, whereas wild-type branch tips began to reorient vertically. These results suggest that *BpLAZY1A* is required for branch gravitropism but is dispensable or functionally redundant in the main stem context.

### *BpLAZY1A* regulates the formation of adaxial tension wood-like tissue in silver birch branches

A tension wood assay was conducted on branches measuring 10-20 cm, 20-30 cm and 30-40 cm in length in WT (V5834) and two *BpLAZY1A RNAi* lines (Fig. S8), using five biological replicates per genotype. We used Astra Blue, a commonly used cationic tension wood stain (e.g. Nugroho *et al*., 2025), that binds to pectins (Kraus *et al*., 1998). Pectins are known to be abundant in the G-layers of tension wood (Mellerowicz and Gorshkova, 2012). We focused on the basal internode, which exhibits the bending phenotype in *BpLAZY1A RNAi* and the *Bplazy1a* mutant (Movie 1; Fig. 2E-F). Our results demonstrate that *BpLAZY1A* mediated adaxial tension wood-like tissue formation is initiated at a very early stage during branch development. In WT plants, adaxial tension wood-like tissue was already evident in young 10-20 cm long branches. Across all developmental stages, adaxial tension wood was observed in 86 % of WT sections. In contrast, 73% of sections from *BpLAZY1 RNAi* line 1 exhibited abnormal tension wood formation, with tension wood either localized to the abaxial side or distributed around the xylem without a clear adaxial polarity. *BpLAZY1 RNAi* line 4 showed a similar phenotype, with 66% of sections displaying abnormal tension wood formation (Fig. 3A-C).

**Figure 3.**
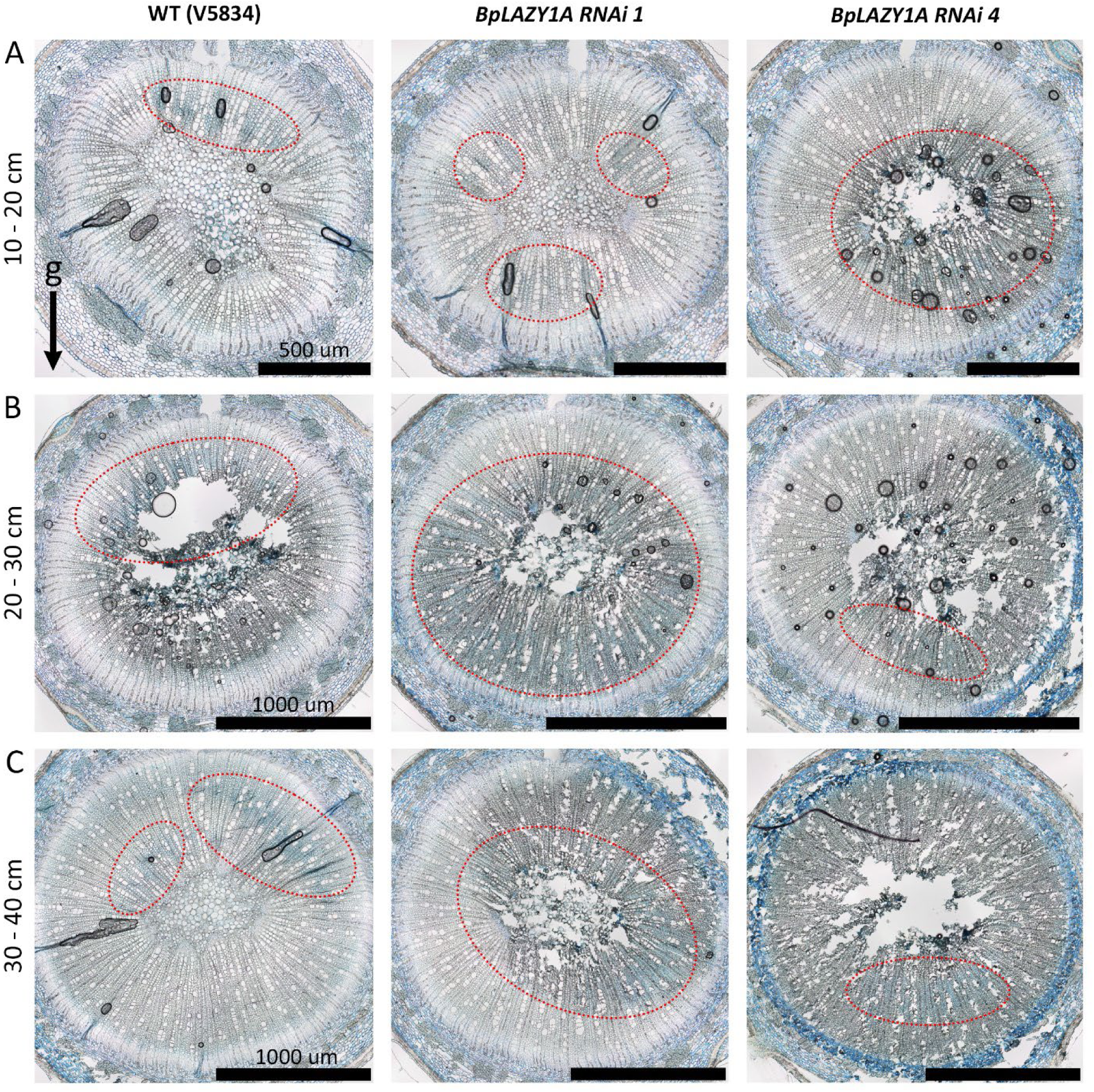
*BpLAZYlA* promotes adaxial tension wood formation in silver birch branches. (A) Adaxial tension wood is evident in young (10-20 cm long) WT branches, whereas tension wood deposition is abnormal in the *BpLAZYl RNAi* lines. (B-C) A similar pattern is observed in older branches, which were 20-30 cm and 30-40 cm in length. In all images, gravity is oriented downward. Red dashed line indicates tension wood localisation. Resin embedded sections (5 µm thick) were stained with 2% Astra Blue in 2% glacial acetic acid.

### *BpLAZY1A* mediates transcriptional polarity in silver birch branches

We investigated the *BpLAZY1A*-mediated transcriptional polarity along the branch axis by conducting RNA sequencing on adaxial and abaxial branch fractions. We analysed adaxial and abaxial tissues from V5834 and *BpLAZY1A RNAi* line 1 at three positions along the branch: a 10 cm apical segment, the third basal internode (mid branch), and the first basal internode (branch base). To reduce inter branch variability, each sample comprised pooled fractions from two adjacent branches per tree (Fig. 4A–B).

**Figure 4.**
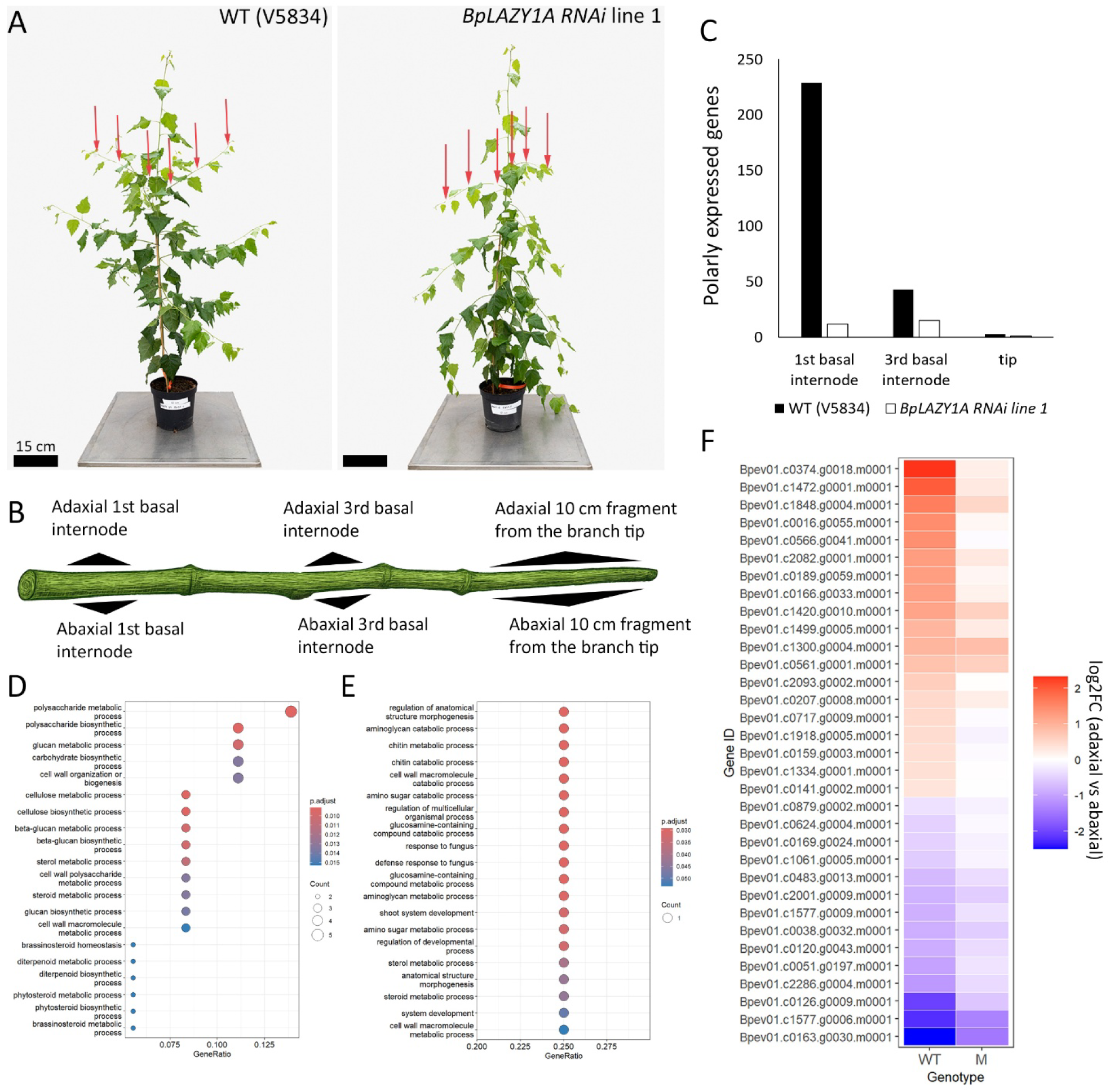
*BpLAZYlA* regulates polar expression of cell wall- and auxin-related genes. (A) Arrows indicate the positions sampled for bulk RNA-seq in WT {VS834) and *8pLAZY1A RNAi* line 1 (B) Schematic of a branch illustrating the sampling positions {not in scale). {C) Bar plot showing the reduced number of polarly expressed genes in *BpLAZY1A RNAi* line 1 compared with WT. {D) GO enrichment analyses highlighting polarized expression of carbohydrate metabolic process related genes in the WT. {E) The polarity in GO enrichment is lacking in *BpLAZY1A RNAi* line 1. {F) Heatmap of the cell wall- and auxin-related genes, which display strong adaxial vs abaxial expression in WT and reduced polarity in *BpLAZYlA RNAi* line 1. Red indicates stronger adaxial expression, blue stronger abaxial expression.

Transcriptional polarity was assessed across all positions within the genotypes. Polarly expressed genes (PEGs) were defined as those exhibiting significant differential expression (adjusted p-value < 0.1; FC 0). We observed a pronounced increase in transcriptional polarity in the wild-type basal internode with 229 PEGs (Fig. 4C; *SI Appendix 1)*. In contrast, the *BpLAZY1A RNAi* exhibited a drastic reduction in polarity with only 12 PEGs (Fig. 4C; *SI Appendix 2*). A similar trend, although with a much smaller difference, was observed in the mid-branch fractions: V5834 and *BpLAZY1A RNAi* had 43 and 15 PEGs, respectively (Fig. 4C; *SI Appendix 3-4*). Transcriptional polarity was virtually absent in apical fractions. V5834 displayed three PEGs and the *BpLAZY1A RNAi* only one (Fig. 4C, *SI Appendix 5-6*). The observed reduction in transcriptional polarity in the mid and the apical fractions may be due to technical constraints. While the greater thickness of the basal fragments allows for precise partitioning into adaxial and abaxial samples, the apical region exhibits higher variability in its orientation relative to the gravitational vector, complicating consistent tissue sampling across multiple samples. Consequently, subsequent analyses were focused on the basal internode, as this region consistently exhibited the bending phenotype in both the *Bplazy1a* segregating population mutant and in the *BpLAZY1A RNAi* (Movie 1, Fig. 2E), adaxial tension wood-like tissue formation in the WT (Fig. 3) and the most robust transcriptional polarity in the WT, which was reduced in the *BpLAZY1A RNAi* (Fig. 4C).

In the basal adaxial fraction, 147 genes were upregulated relative to the abaxial side in the WT, compared to only 12 genes in the *BpLAZY1A RNAi* (*SI Appendix 1-2)*. Six of these PEGs were shared between genotypes, suggesting that a subset of transcriptional polarity is independent of the *BpLAZY1A* function. In the abaxial fraction, 82 genes showed upregulated expression in the WT, whereas no genes were significantly upregulated in the *BpLAZY1A RNAi* (*SI Appendix 1-2*). Taken together, these results demonstrate that the *BpLAZY1A* pathway is a key mediator of transcriptional polarity in silver birch branches.

### *BpLAZY1A* drives polar expression of auxin- and cell wall-related genes

To further characterize the functional implications of this transcriptional polarity, we performed Gene Ontology (GO) enrichment analysis. In V5834 basal internode, polysaccharide metabolic process emerged as the most significantly enriched biological process (Fig. 4D). This finding aligns with histochemical evidence of adaxial tension wood-like tissue formation in wild-type plants, a developmental feature which was irregular in the *BpLAZY1A RNAi* lines (Fig. 3). Furthermore, the strong reduction in PEGs within the *BpLAZY1A RNAi* line 1 was reflected in its markedly limited GO enrichment profile (Fig. 4E).

Based on the assumed *BpLAZY1A* mediated auxin polarity, formation of adaxial tension wood-like tissue in the WT (Fig. 3), and the enrichment of polysaccharide metabolism related processes in the WT (Fig. 4D), we focused our subsequent analysis on auxin- and cell wall-related genes within the basal branch fragments. We manually curated PEGs based on the functional annotations of their closest *Arabidopsis* homologs, excluding genes of unknown function and those that had polar expression in the *BpLAZY1A RNAi*. This filtering yielded a subset of three auxin-related and 30 cell wall-related genes that were polarly expressed in the WT but displayed reduced polarity in the *BpLAZY1A RNAi* (Fig. 4F, *SI Appendix 1*).

On the branch abaxial side, *BpLAZY1A* promotes the expression of two *SAUR-LIKE* genes, which may mediate cell elongation in cortical tissues via the acid growth pathway, consistent with known *SAUR* functions (Stortenbeker & Bemer, 2018; Spartz *et al*., 2014). Additionally, a birch orthologue of *AtIAA29* was upregulated on the abaxial side (*SI Appendix 1*). These results are in line with Kohler *et al*., (2024), who displayed that auxin related genes are more expressed in the abaxial side in the wild-type peach tree branch. Among the 30 cell wall-related PEGs, the largest functional category was associated with pectin metabolism (12 genes). Other identified categories included mannan metabolism (5 genes), cellulose biosynthesis (2 genes), xylan metabolism (2 genes), arabinogalactan metabolism (2 genes), and lignin metabolism (1 gene), alongside one established tension wood marker gene *BpCOBRA-LIKE4*. Additionally, five genes belonged to broader families with currently uncharacterized specific functions in carbohydrate metabolism (Fig. 5A, *SI Appendix 1*).

**Figure 5.**
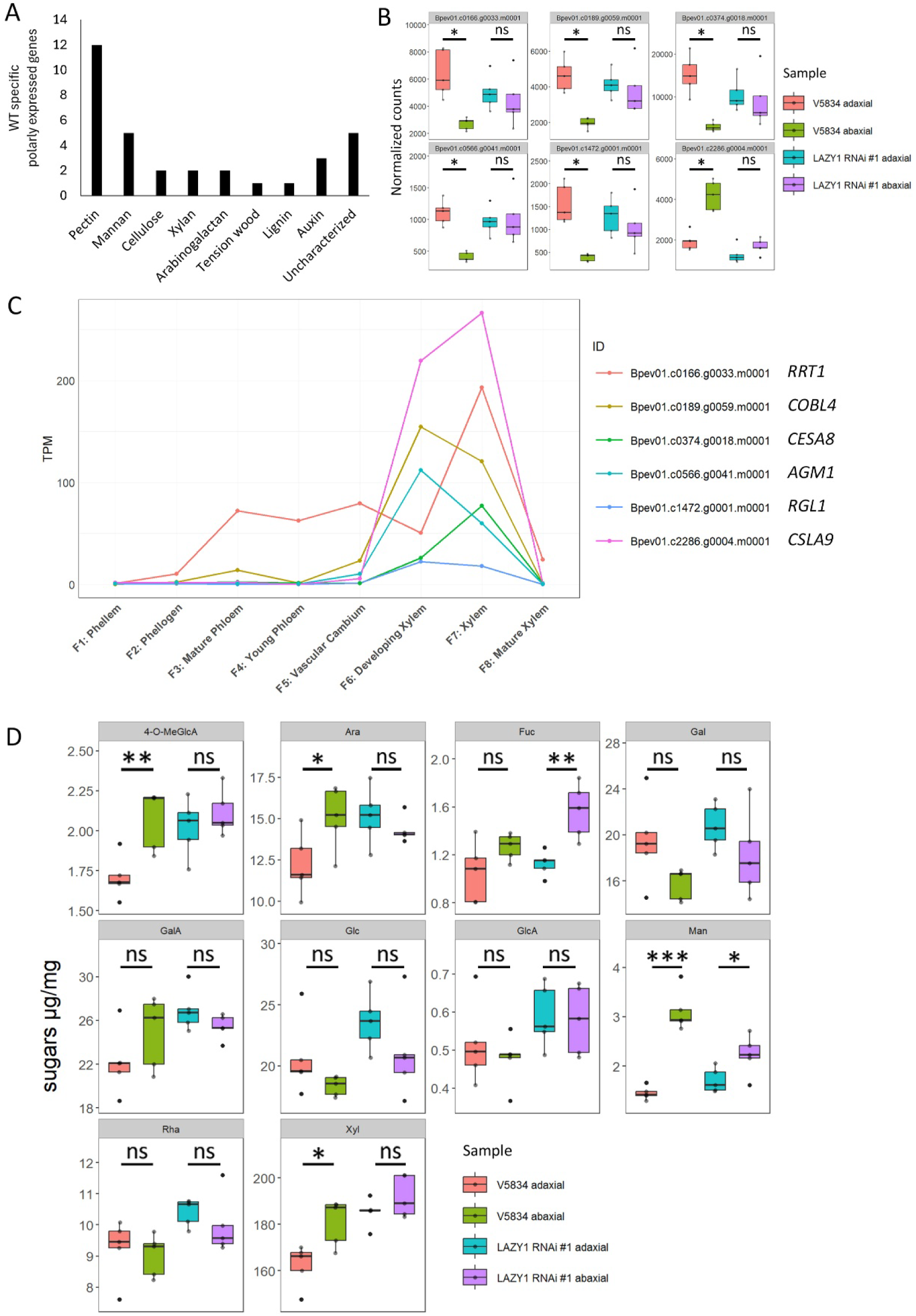
The core cell wall remodeling module of the *BpLAZYlA* pathway. (A) Functional distribution of manually categorized cell wall- and auxin-related genes. These genes exhibit WT specific expression, with polar expression patterns being reduced in *BpLAZYlA RNAi.* Note the enrichment of pectin-related genes within this group. (B) Expression profiles of core components in the *BpLAZYlA-mediated* cell wall remodeling pathway. These genes display strong polarity in the WT (p-adj < 0.001, log,FC > 1, base mean> 1000), which islost in the *BpLAZYlA RNAi.* (C) Tissue-specific transcriptomic profiles from the V5834 main stem. Core components *(RRTl, COBL4, CESAB, AGMl, RGLl, and CSLA9)* show peak expression levels in the xylem and developing xylem. (D) Alcohol Insoluble Residue (AIR) analysis shows statistically significant adaxial-abaxial polarity in WT in 4-O-MeGlcA, Ara and Xyl. Abbreviations: **4-0-MeGlcA, methylated glucuronic acid; Ara, arabinose; Fuc, fucose; Gal, galactose; GalA galacturonic acid; Glc, glucose; GlcA, glucuronic** acid; Man, mannose; Rha, rhamnose; Xyl, xylose. ns; not significant; •; p<0.05; ** a p < 0.01; *** a p < 0.001(Student’s I-test).

Applying a more stringent filtering criteria (log2FC >1 and base mean >1000) narrowed the gene list down to six *BpLAZY1A* dependent branch polarity markers (*SI Appendix 1*). Notably, *BpCOBRA-LIKE4* (Bpev01.c0189.g0059.m0001), the closest birch orthologue to *Populus trichocarpa PtrCOB3*, which is essential for tension wood formation in the poplar main stem (Xu *et al*., 2024; Hsieh *et al*., 2025), exhibited polar expression, with significantly higher transcript abundance on the adaxial side of the WT branches. This polarity was abolished in the *BpLAZY1A RNAi* (Fig. 5B). Four additional genes displayed adaxial-biased expression in V5834: *RHAMNOGALACTURONAN LYASE* (Bpev01.c1472.g0001.m0001), *RG-I RHAMNOSYLTRANSFERASE* (Bpev01.c0166.g0033.m0001), *CELLULOSE SYNTHASE* (Bpev01.c0374.g0018.m0001), and *ARABINOGALACTAN METHYLESTERASE* (Bpev01.c0566.g0041.m0001) (Fig. 5B). RT-qPCR assay confirmed that *BpIAA29* and *BpSAUR-LIKE* displayed abaxially biased polar expression in V5834 and non-polar expression in *BpLAZY1A RNAi* (Fig. S9A-B).

The closest *Arabidopsis* ortholog of the birch *RHAMNOGALACTURONAN LYASE* (Bpev01.c1472.g0001.m0001) is *RGL1* (AT2G22620), which degrades rhamnogalacturonan-I (RG-I) by cleaving the glycosidic linkage between rhamnose and galacturonic acid residues (Min *et al*., 2024). Its poplar ortholog has been shown to reduce fiber cell-to-cell adhesion (Yang *et al*., 2020). The *RG-I RHAMNOSYLTRANSFERASE* (Bpev01.c0166.g0033.m0001) is orthologous to *RRT1* (AT5G15740), a Golgi-localized enzyme that catalyses the transfer of rhamnose to the RG-I backbone (Takenaka *et al*., 2018). The birch *CELLULOSE SYNTHASE* (Bpev01.c0374.g0018.m0001) is orthologous to *CESA8* (AT4G18780), which regulates crystallinity, crystal width, and longitudinal deposition of cellulose microfibrils in *Arabidopsis* (Hill *et al*., 2025). Notably, these structural qualities of cellulose are defining features of tension wood cellulose (Clair *et al*., 2011; Mellerowicz & Gorshkova, 2012).

The closest *Arabidopsis* orthologue of the birch *ARABINOGALACTAN METHYLESTERASE* (Bpev01.c0566.g0041.m0001) is *AGM1* (AT1G27930), which methylates glucuronic acid (GlcA) residues at the termini of beta-1,6-galactan sidechains and within the beta-1,3-galactan backbone of type II arabinogalactans (AGs) (Temple *et al*., 2019). These highly branched polysaccharides are covalently linked to hydroxyproline residues of plant cell wall proteins to form arabinogalactan proteins (AGPs), a diverse class of cell-surface proteoglycans ubiquitous across the plant kingdom (Showalter, 2001; Temple *et al*., 2019). While the functionality of GlcA methylation of AGs remains to be elucidated, a T-DNA knockout lines of a β-1,6-glucuronyltransferase, an enzyme that transfers GlcA to the AG backbone, exhibit increased elongation rates in *Arabidopsis* roots and hypocotyls (Knoch *et al*., 2013; Temple *et al*., 2019).

Among the genes highlighted by this filtering, only one gene, *BETA-MANNAN SYNTHASE* (Bpev01.c2286.g0004.m0001) exhibited higher transcript abundance on the abaxial side (Fig. 5B). Its closest *Arabidopsis* orthologue is *CELLULOSE SYNTHASE-LIKE A9* (At5g03760), which synthesizes the β-1,4-mannan backbone of galactomannan (Davis *et al*., 2010). β-mannans are structurally varied polysaccharides that associate with cellulose microfibrils. Glucomannan backbones develop diverse patterns of galactosyl substitution that vary with plant species and developmental stage. Yoshimi *et al*. (2025) demonstrated that the extent of galactosylation plays a key role in modulating glucomannan solubility and its interaction with cellulose. Our data is in line with (Andersson-Gunnerås *et al*., 2005) who displayed that mannan biosynthesis related genes in poplar tension wood are downregulated compared with normal wood in poplar main stem. However, reduced level of mannans in poplar main stem does not have an impact on tree height, radial growth or total biomass (Guevara-Rozo *et al*., 2025). Reduced quantities of β-mannans might therefore be important and specific for the function of tension wood.

The six cell wall related genes that passed the stringent filtering criteria (log2FC > 1; base mean > 1000) exhibit peak expression in developing xylem or mature xylem tissues in silver birch (Fig. 5C; data from Alonso-Serra *et al*., 2019). Although tissue-specific transcriptomic data are not available for silver birch branches, the main-stem dataset (Alonso-Serra *et al*., 2019) provides a relevant reference for establishing the secondary xylem-specific expression and putative roles of these genes in tension wood formation.

### Monosaccharide composition analysis

Following our transcriptomic data that points to the importance of the *BpLAZY1A* pathway in cell wall remodelling, we proceeded with a monosaccharide analysis. Tension wood is typically characterized by elevated levels of cellulose, xyloglucan, rhamnogalacturonan I (RG-I), de-arabinosylated RG-I, galactans, arabinogalactans, and type II arabinogalactan proteins (AGPs), alongside reduced xylan and mannan relative to normal wood (Mellerowicz & Gorshkova, 2012). No clear enrichment of these tension wood markers was detected in the adaxial branch samples of the WT. In WT, xylose and methylated glucuronic acid levels were lower on the adaxial side, whereas this difference was not statistically significant in *BpLAZY1A RNAi* line 1. Mannose levels were lower on the adaxial side in both WT and *BpLAZY1A RNAi* line 1. Fucose levels were higher on the abaxial side in *BpLAZY1A RNAi* line 1, but no significant difference was detected in WT. Together, these results likely reflect limitations in sample resolution, as bulk fractionation combined multiple cell types within adaxial and abaxial tissues. Such limitations may be negligible in the decapitation context, where polarity differences became extremely pronounced (Fig. S7A-B), but becomes critical in branches, where differences are much more subtle (Fig. 3A-C). Consequently, spatial approaches such as Raman spectroscopy will be essential to resolve differences in wood chemical composition between adaxial and abaxial tissues in the branch context.

## Discussion

While *LAZY* mediated gravitropism has been extensively characterized in *Arabidopsis* roots, our results extend the gravitropism model to the shoot system of a woody species undergoing secondary growth, where polarity must be coordinated across multiple tissue types. We identify *BpLAZY1A* as a central mediator of transcriptional polarity and driver of adaxial tension wood-like tissue formation in silver birch (*Betula pendula*) branches. In contrast to previous studies (Wilson and Archer, 1983; Kohler *et al.,* 2024), we were able to regularly detect adaxial tension wood-like tissue in young WT branches. The contrast of tension wood vs opposite wood, however, becomes much stronger after decapitation context, which we assume, is when the uppermost branches acquire the main stem identity. The pronounced reduction in PEGs in *BpLAZY1A RNAi* demonstrates that *BpLAZY1A* is required for translating the assumed auxin asymmetry into coordinated tissue dependent transcriptional programs. Auxin related genes are enriched on the abaxial side, whereas cell wall modifying genes are preferentially expressed adaxially. Our data supports a model in which auxin simultaneously promotes abaxial cell elongation while driving tension wood-like differentiation in the adaxial xylem (Fig. 6). The adaxial enrichment of pectin associated genes, together with the polar expression of *BpCOBRA-LIKE4* and *BpCESA8*, indicates that *BpLAZY1A* dependent polarity modulates both pectin deposition and cellulose microfibril organization, which likely affect the mechanical properties of the branch specific tension wood-like tissue. Our on-going work studies whether there is only a quantitative difference in branch specific tension wood or is it also qualitative. However, loss of transcriptional polarity in *BpLAZY1A RNAi* correlates with abnormal tension wood-like tissue formation and altered branch posture, underscoring the functional link between the transcriptional asymmetry and the biomechanical output.

**Figure 6.**
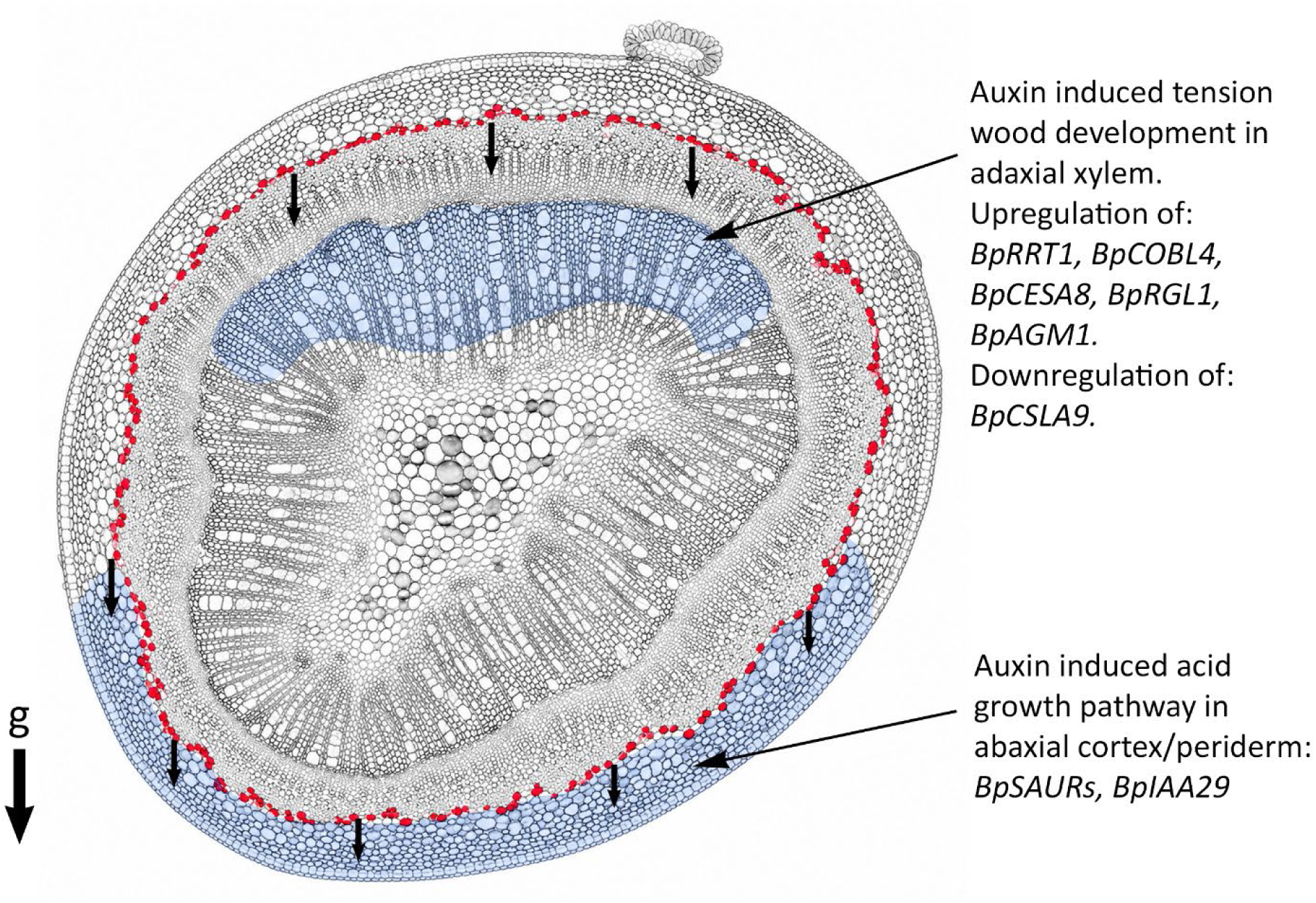
Model on *BplAZYlA* mediated adaxial-abaxial polarity in silver birch branches. The *BpLAZYlA* complex mediates gravity directed auxin efflux from the starch sheath (red cells) via PIN recruitment and activation. This establishes an asymmetric auxin gradient across the vascular cambium, resulting in differential auxin concentrations between the adaxial and abaxial sides. On the adaxial side, this hormonal polarity triggers tension wood development (generating a pulling force). This mechanism involves the upregulation of the tension wood marker *BpCOBL4,* pectin metabolism genes *(BpRRTl, BpRGLl),* highly crystalline cellulose deposition along the longitudinal cell axis *(BpCESAB),* and glucuronic acid methylation on arabinogalactan *(BpAGMl),* alongside the downregulation of beta-mannan biosynthesis *(BpCSLA9).* Conversely, on the abaxial side, the *BpLAZYlA* pathway activates the acid growth pathway in the cortex/periderm via *BpSAURs,* driving enhanced abaxial expansion (generating a pushing force).

Taken together, our findings support a model in which *BpLAZY1A* mediates adaxial tension wood-like tissue formation in silver birch branches and possibly drives abaxial cell expansion in through the acid growth pathway. This model provides a framework for understanding how plagiotropic branch growth is maintained in trees and how this pathway could be tuned for various agricultural applications.

## Materials and methods

See supplementary file

## Supporting information

Movie 1 - straight

Movie 2 - tilted

Movie 3 - decapitated

SI 7 silver birch transformation protocol

## Acknowledgements

We thank Junko Takahashi and Sonja Viljamaa at the Biopolymer Analytical Platform (BAP) at UPSC/SLU, supported by Bio4Energy for the pyrolysis-GC/MS and the monosaccharide composition analysis. We also thank the greenhouse facility staff at the University of Helsinki Viikki campus for constant support.

The authors acknowledge the use of ChatGPT-5.5 (OpenAI), Gemini 3.1 Pro (Google) and Claude Sonnet 4.6 (Anthropic) for language and R code refinement.

## SUPPLEMENTARY FIGURES

**Figure S1.**
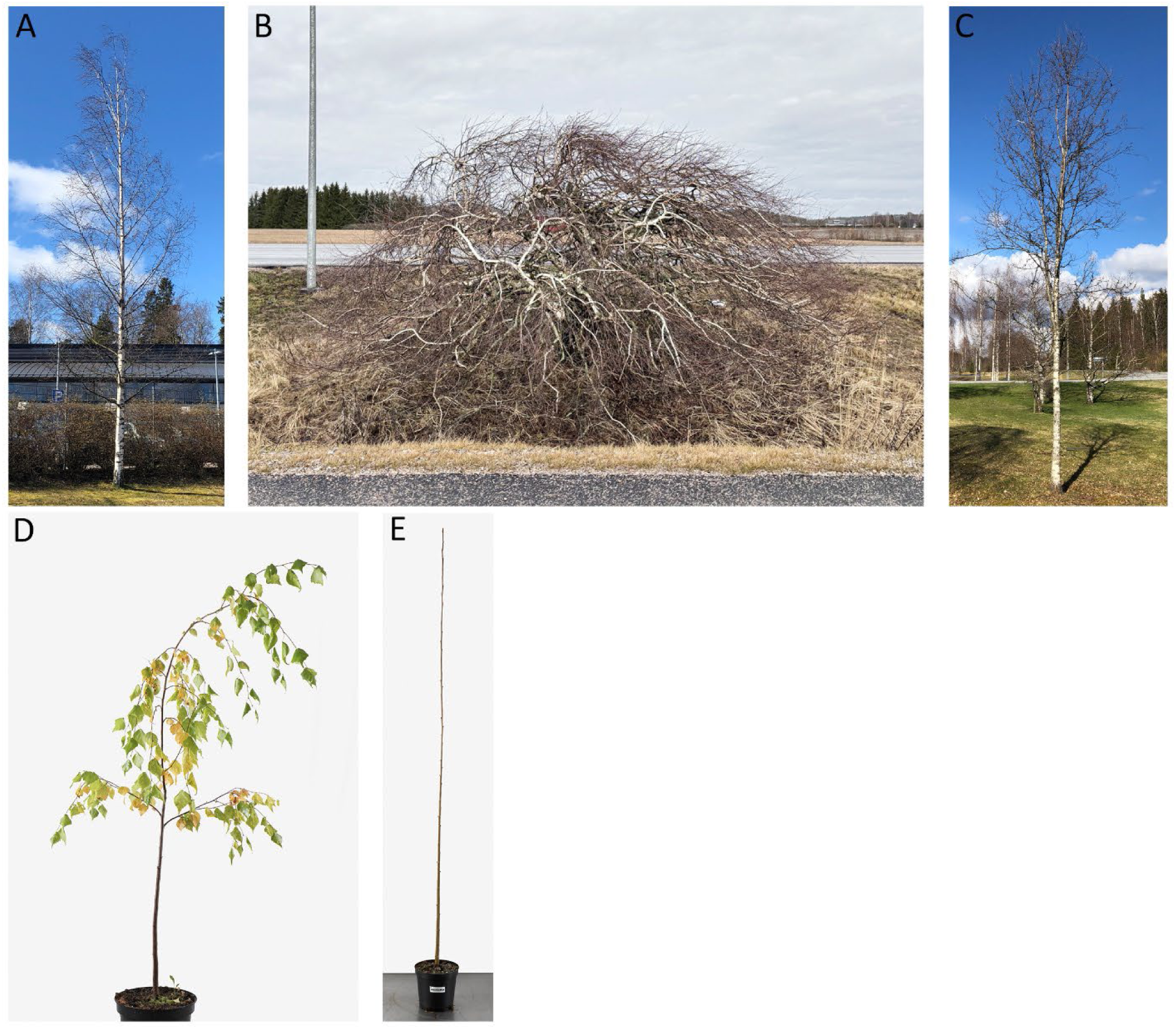
Phenotypic variation and inheritance patterns of various birch shoot system mutants. (A) Non-pendulous *Betu!a pendula.* (B) Bushy *B. pendula.* Likely recessive. F, is 100% WT-like. (C) Upright *B. pendula.* Likely recessive. F1 is 100% WT-like. (D) Semi-weeping *B. pendu/a.* Inheritance pattern unknown. (E) Stunted branches in a tetraploid *B. pubescens.* Likely semi-dominant. F, is segregating 9:4 (WT:M) at 3-month-old age in a 52 plant population. A larger population and extended phenotyping period is required to confirm the branch phenotype and segregation pattern in putative F, mutants.

**Figure S2.**
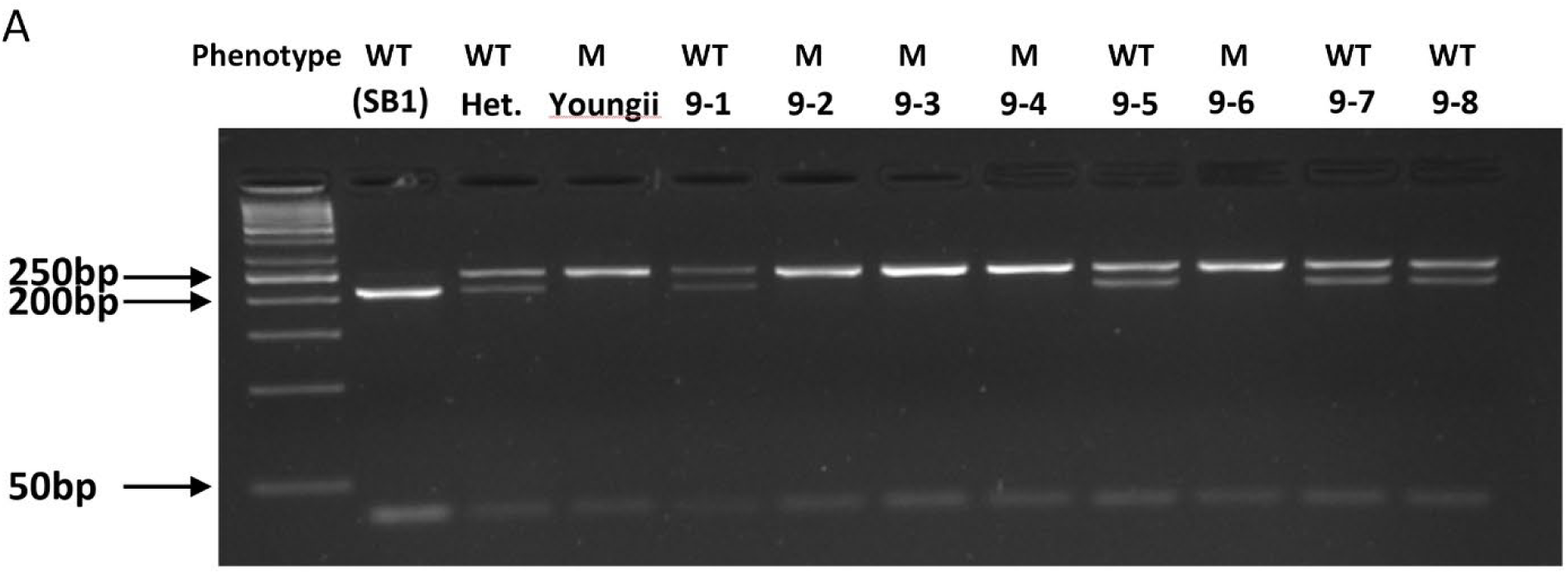
Restriction enzyme digestion assay confirms the SNP in the *BpLAZYlA* gene in the segregating population. (A) A 293 bp gDNA fragment of the *BpLAZYlA* gene was amplified by PCR and digested with Dpnll. The expected fragment sizes were 37 bp, 219 bp, and 256 bp for heterozygotes; and 37 bp and 256 bp for homozygote mutants. A subset of the genotyped segregating population individuals is displayed here.

**Figure S3.**
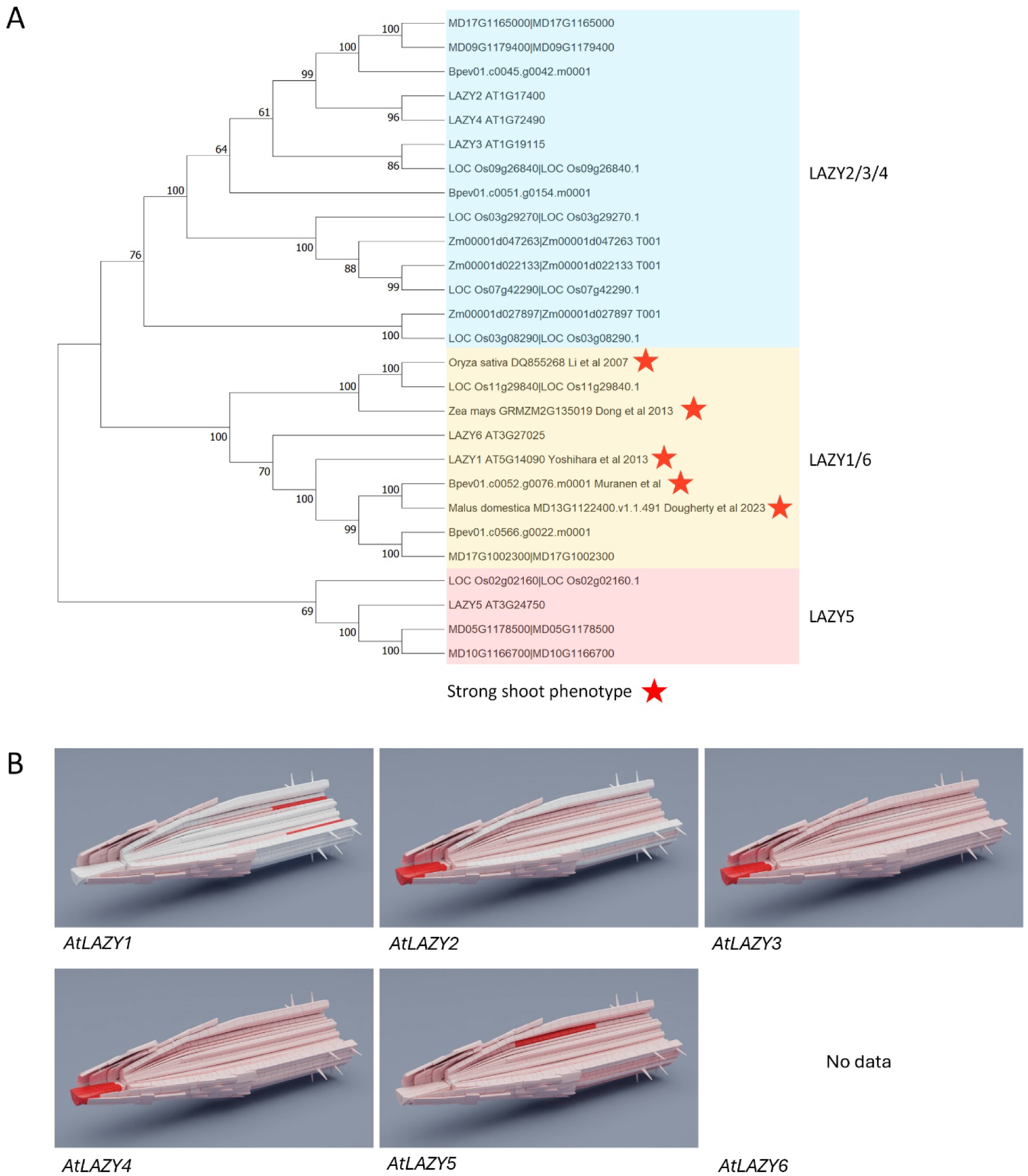
Mutations in the *LAZYl/6* clade underlie strong shoot phenotypes across multiple species. (A) UPGMA inferred phylogenetic relationships among the *LAZY* gene family members in silver birch *(Betula pendu/a),* Arabidopsis, rice *(Oryza sativa),* maize *(Zea mays),* and apple tree *(Ma/us domestica).* The values indicated on the branches represent the bootstrap values for 1000 replicates. Reported mutations associated with severe shoot phenotypes consistently fall within the *LAZYl/6* clade, indicating that *LAZYl/6* clade functions primarily in shoot gravitropism across species. The role of *AtLAZY6* in gravitropism is currently unknown. (B) scRNA-seq data of the *LAZY* gene family in the *Arabidopsis* root apex display that *AtLAZYl* is the only member which is strongly expressed in the endodermis (shoot gravitropism), while *AtLAZY2-4* show strong expression in the columella cells (root gravitropism). *AtLAZY5* exhibits its highest expression in the cortex. Expression data was not available for *AtLAZY6* in this dataset. These patterns support that *LAZYl* plays a central role in shoot gravitropism.

**Figure S4.**
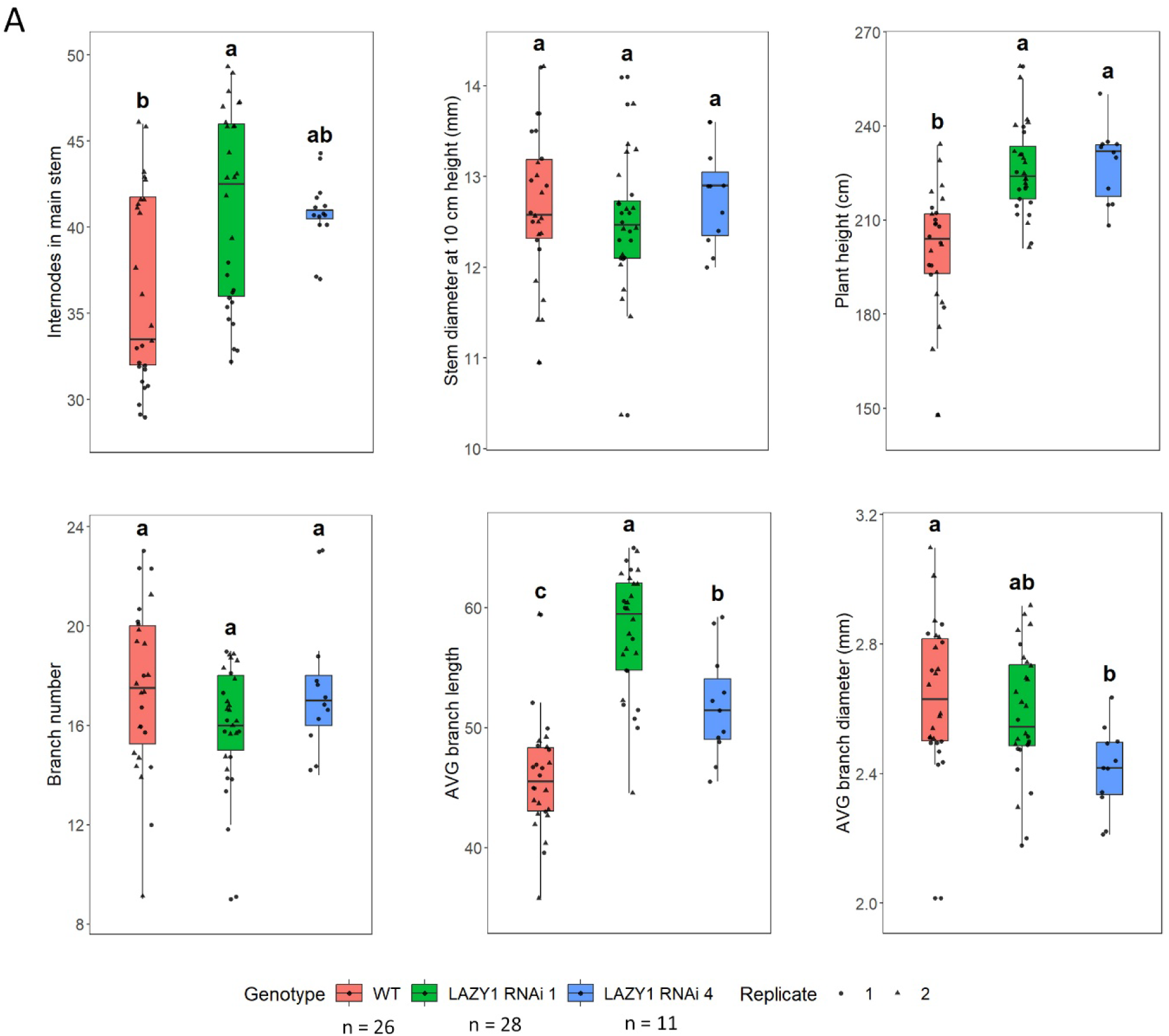
*BpLAZYlA RNAi* plants grow taller and dispaly increased branch length. Plant swere phenotyped three motnhs after potting. *BpLAZYlA* pathway affects apical growth. *BpLAZYl RNAi* plants grow taller and they grow longer branches. Statistical test: One-way ANOVA followed by Tukey HSD. Letters indicate significance difference (p-value <0,05). Data comes from 26 (WT), 28 *(BpLAZYlA RNAi 1)* and 11 *(BpLAZYlA RNAi 4)* biological replicates from two separate experiments, except for *BpLAZYlA RNAi 4* that was phenotyped only once.

**Figure S5.**
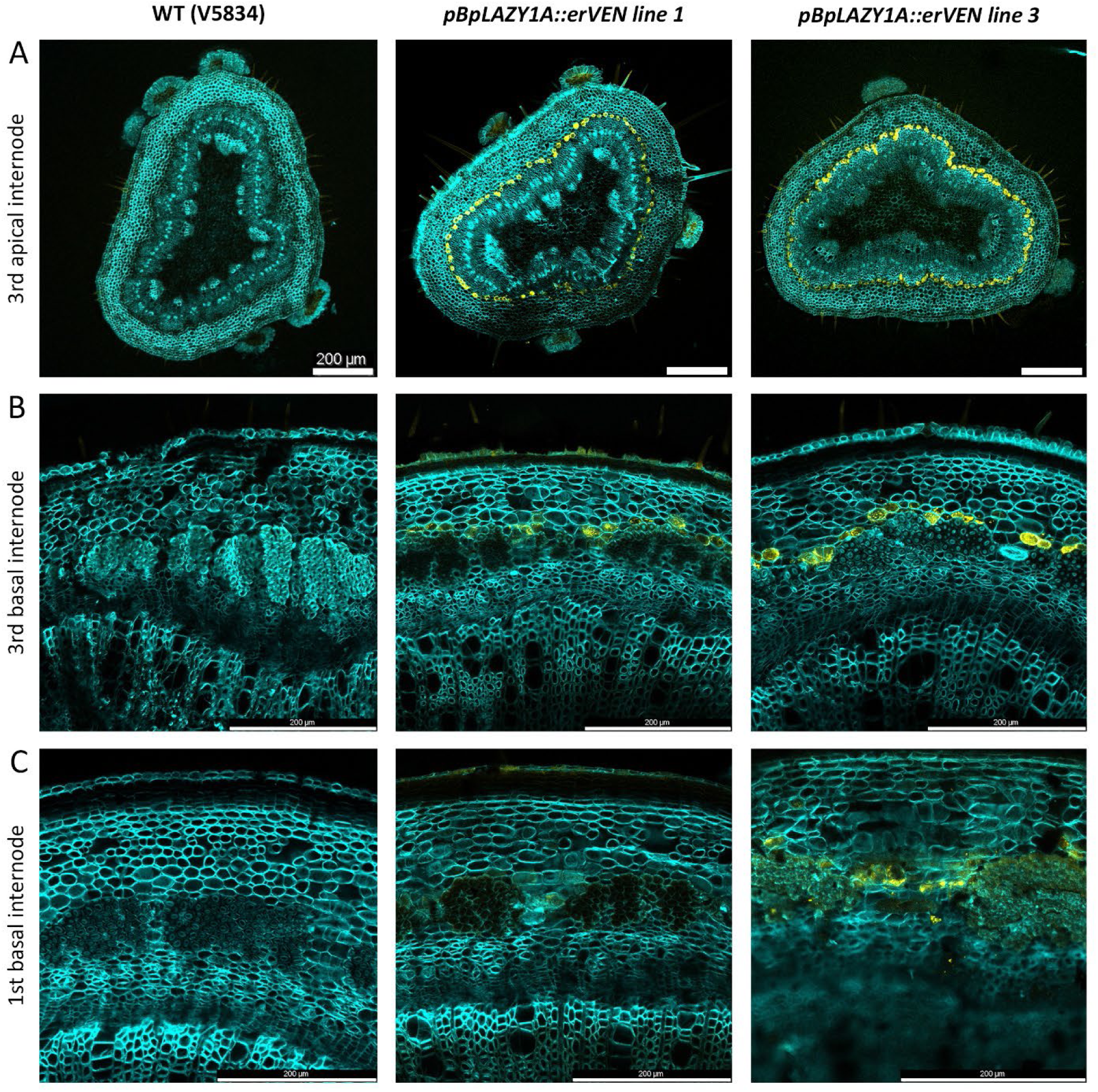
*BpLAZYlA* is expressed in the starch sheath throughout the branch axis. (A) The *BpLAZYlA::erVen* reporter shows that the promoter is active in the starch sheath in the 3rd apical internode at the branch tip. (B) Reporter activity is also observed in the third basal internode, corresponding the middle region of the branch. (C) Reporter activity is additionally detected in the basal internode of the branch. Wild type is included as an autofluorescence control.

**Figure S6.**
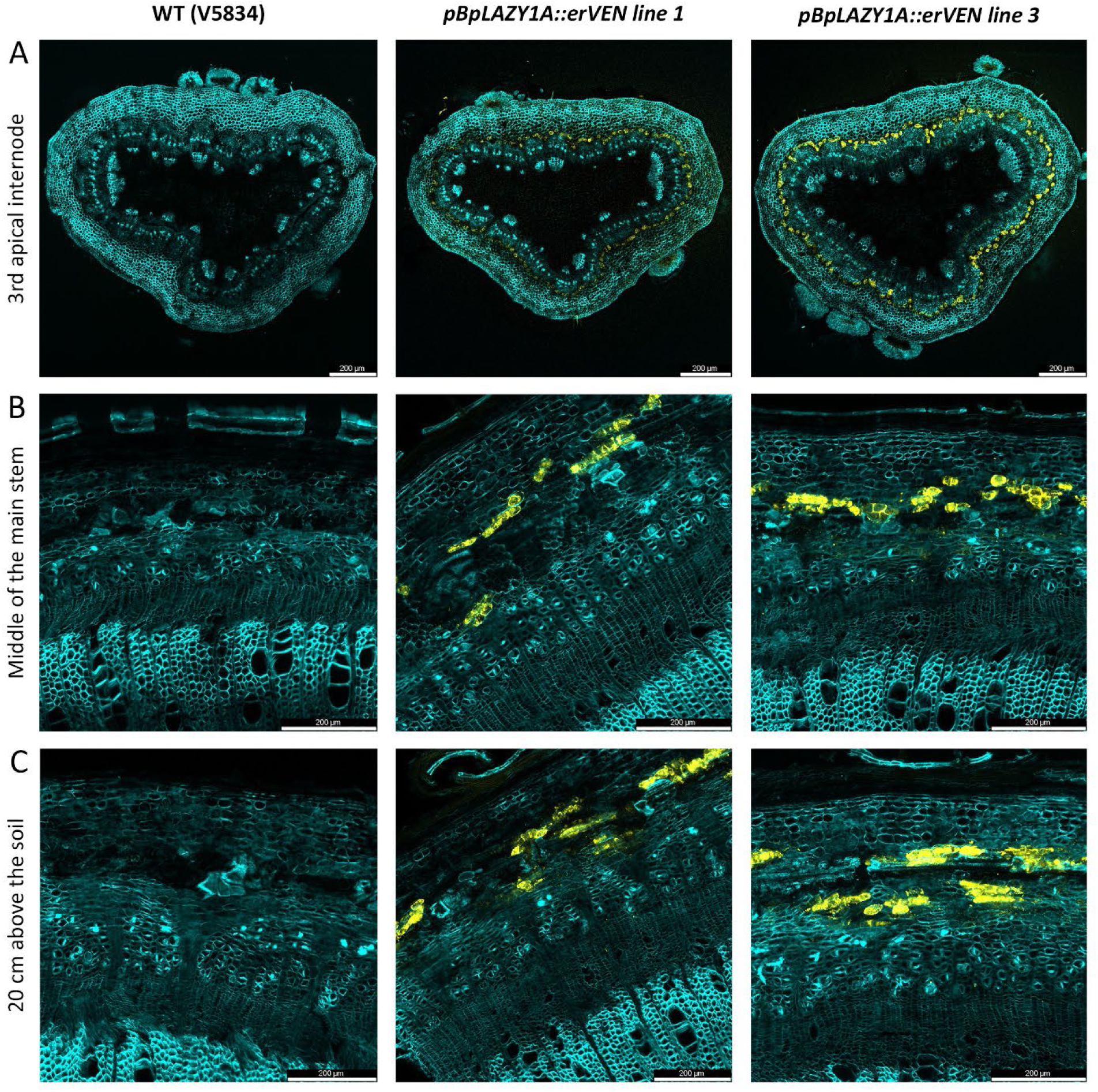
*BplAZYlA* is expressed in the starch sheath throughout the main stem. (A) The *BpLAZYlA::erVen* reporter indicates promoter activity in the starch sheath of the third apical internode at the branch tip. (B) **In** the middle region of the main stem, reporter activity is detected in the starch sheath and occasionally in phloem cells adjacent to the phloem fiber bundles. (C) In main stem regions 20 cm above the soil, reporter activity is observed in the starch sheath and occasionally in phloem cells adjacent to the phloem fiber bundles.

**Figure S7.**
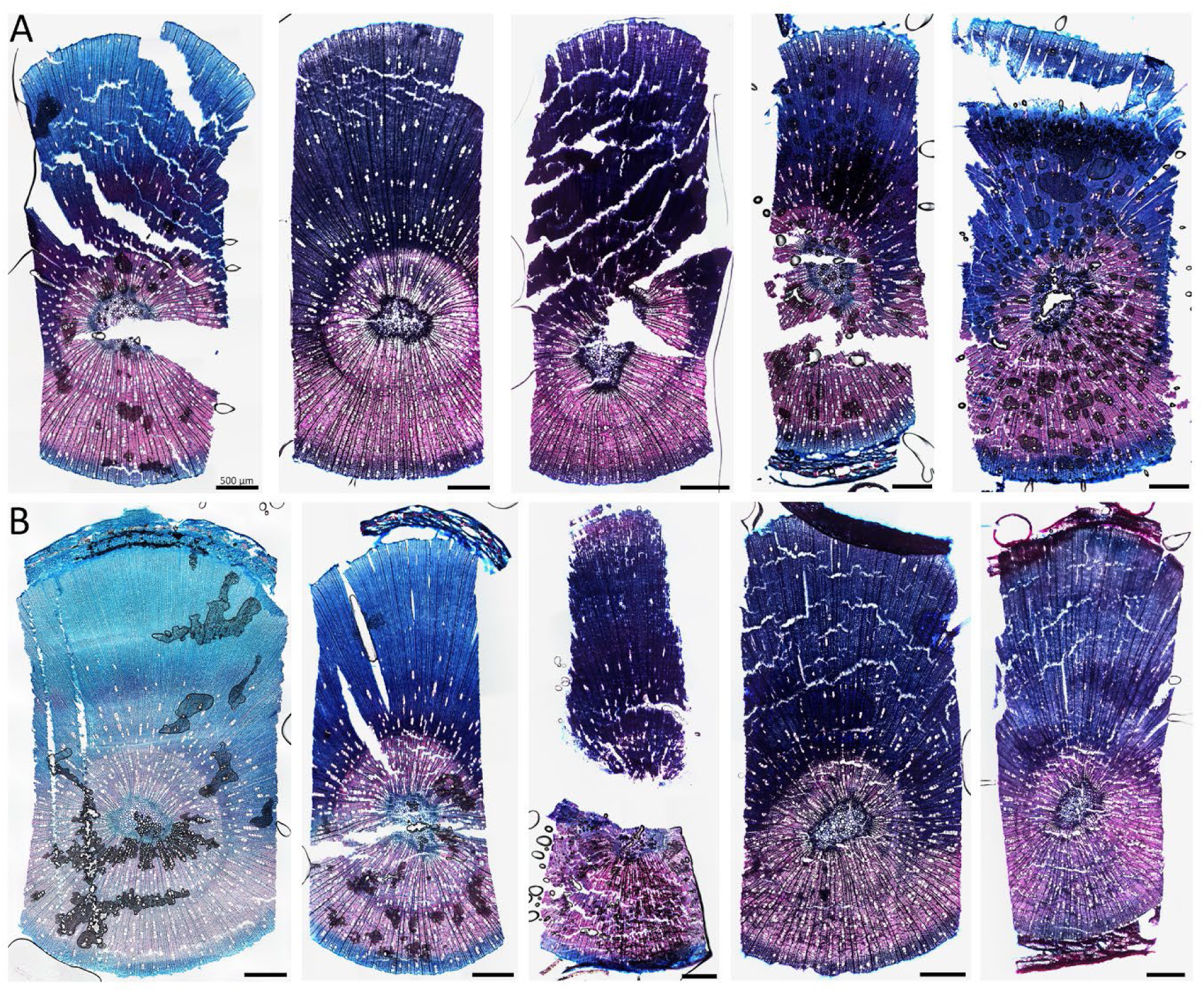
Decapitation induces extensive tension wood formation in WT and *BplAZYlA RNAi*. (A) Extensive and abrupt tension wood formation is observed four weeks after decapitation in all WT cryo-sections (5/5) from the branch base. (B) A similar response is observed in all *BplAZYlA RNAi* samples (5/5). Sections are stained with Safranin O and Alcian Blue.

**Figure S8.**
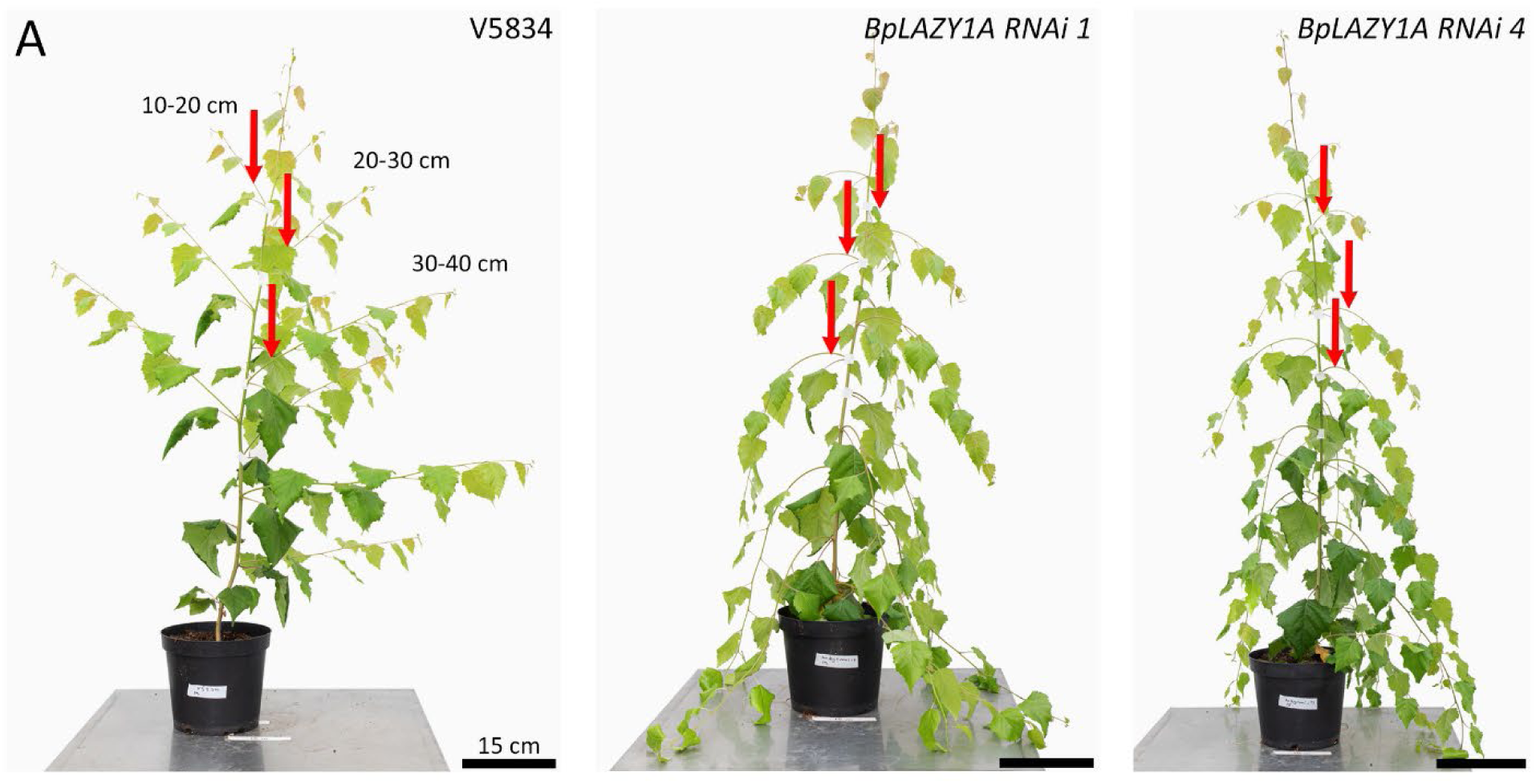
Tension wood sampling positions by developmental stage. (A) Tissue samples were collected from the branch bases of WT (VS834), *BpLAZYlA RNAi* line **1,** and line **4** at branch lengths of 10-20 cm, 20-30 cm, 30-40 cm.

**Figure S9.**
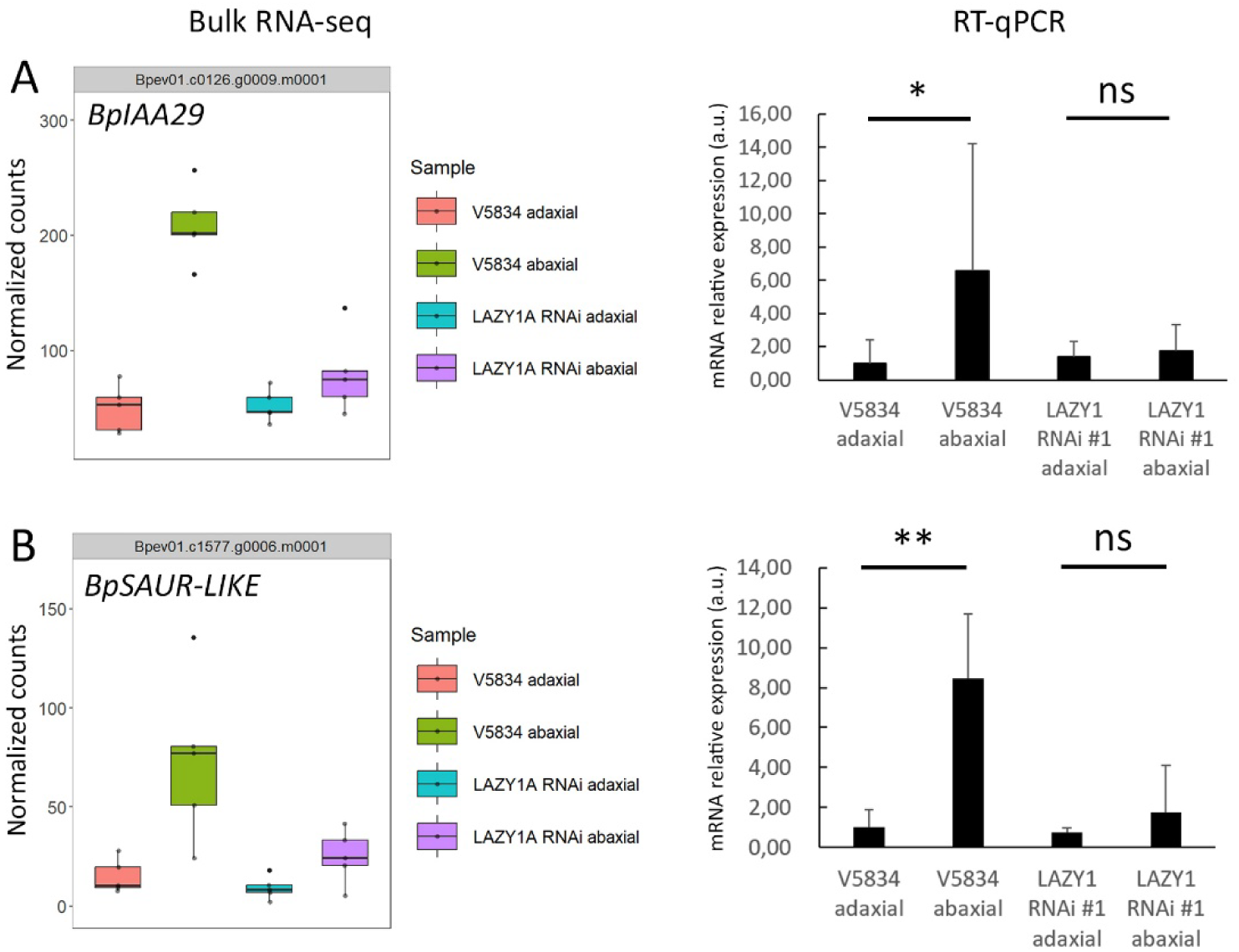
RT-qPCR validation of bulk RNA-seq data. (A) *Bp/AA29* exhibits polar expression in WT (VS834) in bulk RNA-seq, but not in the *BpLAZYlA RNAi.* RT-qPCR confirms polar expression in WT and non-polar expression in *BpLAZYlA RNAi.* (B) *BpSAUR-LIKE* exhibits polar expression in WT in bulk RNA-seq, but not in the *BpLAZYlA RNAi.* RT-qPCR confirms polar expression in WT and non-polar expression in *BpLAZYlA RNAi.* RT-qPCR data are represented as mean ±1 SD. N = 3 biological replicates. ns, not significant; * p < 0.05; ** p < 0.01 (Student’s t-test).

## Notes

### Competing Interest Statement

The authors have declared no competing interest.

