## Supplementary material for "*BpLAZY1A* Mediates Transcriptional Polarity and Drives Adaxial Tension Wood–Like Tissue Formation in Silver Birch (*Betula pendula*) Branches": SI 7 silver birch transformation protocol

Adapted from Keinonen-Mettälä et al. (1998). Comparisons of the efficiency of some promoters in silver birch (Betula pendula). *Plant Cell Reports.* Edited by Sampo Muranen 20/5/2026.

Read protocol carefully before starting. Most of the steps contain pre-tasks. Use appropriate selection for the Agrobacterium strain and for the plant selection.

Expression vector: Vector of interest with kanamycin resistance

Agrobacterium strain: c58 GV3101 pMP90 (Koncz and Schell, 1986) Rif and Gen res.

#### 1. Agro transformation + growing on plates (2 days + PCR)

-Transform competent agro cells with 1 µl of expression vector extracted plasmid solution (100 - 300 ng). Mix with 200 µl SOC medium. Incubate at +28° for 2 h.
-Pipette 5 and 10 µl transformed agro per plate (2 plates is usually enough).
-Grow at +28°C for two days. Store colonies at +4°.

**Pre-task**: Prepare 4 selection plates according to the construct (100 ml)

| | |
|---|---|
| • LB agar | **100 ml** |
| • Rifampicin 100 µg/ml | 200 µl from 50 mg/ml stock |
| • Gentamycin 25 µg/ml | 250 µl from 10 mg/ml stock |
| • Kanamycin 50 mg/L (binary vector) | 25 ul from 100 mg/ml stock |

#### 2. Plant material pre-culture on plates in dark (5 days before transformation)

-Cut in vitro stems (when they are around 8 cm tall) into 4 cm bits.
-Set stem pieces and leaves in pre-cultivation plates
-Seal plates with parafilm
-Incubate on plates (vertically) for 5 days in dark at RT.

**Pre-task**: prepare pre-cultivation plates (15-20 plates).

| **Pre-cultivation media** | **500 ml** |
|---|---|
| • WPM 2,46 g/L | 1,23 g |
| • MES 0,5 g/L | 0,25 g |
| • sucrose 20 g/L | 10 g |
| • Agar 7 g/L | 3,5 g |
| • set pH 5,6 | |
| • Autoclave | |

**Add when making plates**

| | |
|---|---|
| • BAP 1 mg/L | 500 ul from 1mg/ml stock |
| • 2,4-D 2 mg/L | 1000 ul from 1mg/ml stock |

**Prepare these**
-Prepare agro re-suspension culture with PVP and co-culture medium with PVP (step 4).
-Autoclave box of 5 ml tips (step 4).
-Prepare washing medium (step 5).
-Autoclave small sheets of filter paper inside foil (step 6).
-Prepare selection media (step 6).

#### 3. Agro culture overnight

-Transfer 8 separate colonies into 8*5 ml **Agrobacterium suspension media**.
-Incubate at 28°C, 160 rpm, overnight.
-Check construct with Dreamtag PCR (1 ul media in a 25 ul PCR reaction). Remember to vortex before pipetting suspension out of the 5ml tube!

**5 ml agro suspension media**

| | |
|---|---|
| • LB | **5 ml** |
| • Rifampicin 100 µg/ml | 10 µl from 50 mg/ml stock |
| • Gentamycin 25 µg/ml | 12,5 µl from 10 mg/ml stock |
| • Kanamycin 50 mg/L (Bin. vector) | 1,25 ul from 100 mg/ml stock |

#### 4. Co-culture

-Agro suspension culture 100 ml overnight. At 16:00 transfer 5 ul and 10 ul from 5 ml agro suspension culture into **2*100 ml Agrobacterium suspension media. Remember to vortex before collecting!** Collect 2 ml culture for the OD600 reference check the morning after.

**100 ml Agrobacterium suspension media**

| | |
|---|---|
| • LB | 100 ml |
| • Rifampicin 100 µg/ml | 200 µl from 50 mg/ml stock |
| • Gentamycin 25 µg/ml | 250 µl from 10 mg/ml stock |
| • Kanamycin 50 mg/L (Bin. vector) | 25 ul from 100 mg/ml stock |

-Grow overnight at 28°C (to OD600 0.3 **-** 0.4) in shaker 160 rpm.
-Centrifuge in 2 x 50 ml Falcons at 4000g for 10 min at +4°C.

**Agrobacterium re-suspension media with PVP**
-Re-suspend pellet with **Agro re-suspension culture with PVP.**

| **Agro re-suspension media with PVP** | **100 ml** |
|---|---|
| • MS with vitamins 2,14 g/L | 0,214 g |
| • Sucrose 20 g/L | 2 g |
| • MES 0,5 g/L | 0,05 g |
| • PVP 40 10 g/L | 1 g |
| • set pH 5.6 | |
| • Autoclave | |

**Add before use**

- Acetosyringone 20 mg/L — 400 ul from 5mg/ml stock

**Wounding of stem pieces**

-Transfer stem pieces and leaves from pre-culture plates into **ascorbic acid media plate**. Use large rectangular plates. Make few holes in the stems and leaves with a small syringe needle. Move onto a new **ascorbic acid media plate and cut into 2 cm pieces**. Transfer into **agrobacterium re-suspension media with PVP** as you cut.

| **Ascorbic acid media** | **250 ml** |
|---|---|
| • MS with vitamins 2,14 g/L | 0,535 g |
| • Set pH to 5.6 | |
| • Autoclave | |

**Add before use**

- Ascorbic acid ~750 mg/L — 2 ml from 100 mg/ml stock (filter sterilized)

**Co-culture**

-After pierced stem pieces and leaves have been transferred into 2 x 50 ml **Agrobacterium re-suspension media with PVP**, put into a falcon tube rotator for 2 h (slow). Keep tubes inside foil.

-Transfer plant material into **co-culture media with PVP** (change the media daily in the same tube).

-Culture 5 days in dark (foil around tubes) at RT, in a rotator (slow).

| **Co-culture media with PVP** | **100 ml** |
|---|---|
| • MS with vitamins 2,14 g/L | 0,214 g |
| • Sucrose 20 g/L | 2 g |
| • MES 0.5 g/L | 0,05 g |
| • PVP 10 g/L | 1 g |
| • set pH 5.6 | |
| • Autoclave | |

**Add before use**

| | |
|---|---|
| • TDZ 0,5 mg/L | 50 ul from 1mg/ml stock |
| • NAA 0,2 mg/L | 20 ul from 1mg/ml stock |
| • Acetosyringone 10 mg/L | 200 ul from 5mg/ml stock |

#### 5. Washing

- Rinse plant material in 3*100 ml sequential **washing media** (1h 100ml with antibiotics + 1h 100 ml with antibiotics + 1h 100 ml in MS)
- Dry on sterile filter paper

| **Washing media** | **300 ml** |
|---|---|
| • MS with vitamins 2,14 g/L | 0,65 g |
| • MES 0,5 g/L | 0,15 g |
| • Sucrose 20 g/L | 2 g |
| • Set pH to 5.6 | |
| • Autoclave | |

| **Add before use** | **100 ml** |
|---|---|
| • Claforan 500 mg/L | 200 ul from from 250 mg/ml stock |
| • Vancomycin 500 mg/L | 500 ul from 100 mg/ml stock |

#### 6. Transformed plant material on selection and shoot induction plates (months)

-Transfer plant material onto **selection plates.**
-Keep under foil for the first 7 days. Then in low light 50 uE (cover with a thin paper sheet).
-Change Kanamycin C from 25 mg/L to 50 mg/L after two weeks.
-Every two weeks: transfer material onto fresh plates until large (~1 $cm^3$) callus forms. Usually this happens within two months. After callus has formed, use **shoot induction media.**

| **Selection media** | **500 ml** |
|---|---|
| • WPM 2,46 g/L | 1,23 g |
| • MES 0,5 g/L | 0,25 g |
| • sucrose 20 g/L | 10 g |
| • Agar 7 g/L | 3,5 g |
| • set pH 5.6 | |
| • Autoclave | |
| **Add when making plates** | |
| • TDZ 0,5 mg/L | 250 ul from 1 mg/ml stock |
| • NAA 0,2 mg/L | 100 ul from 1 mg/ml stock |
| • Kanamycin 25-50 mg/L (plant selection) | 62,5-125 ul from 100 mg/ml stock |
| • Claforan 250 mg/L | 500 ul from from 250 mg/ml stock |
| • Vancomycin 200 mg/L | 1000 ul from 100 mg/ml stock |

| **Shoot induction plates** | **500 ml** |
|---|---|
| • WPM 2,46 g/L | 1,23 g |
| • MES 0,5 g/L | 0,25 g |
| • sucrose 20 g/L | 10 g |
| • Agar 7 g/L | 3,5 g |
| • set pH 5.6 | |
| • Autoclave | |

**Add when making plates**

- BAP 0,8 mg/L — 400 ul from 1 mg/ml stock
- NAA 0,02 mg/L — 10 ul from 1 mg/ml stock
- GA3 0.5 mg/L — 250 ul from 1 mg/ml stock
- Kanamycin 25 mg/L (plant selection) — 125 ul from 100 mg/ml stock
- No kanamycin after shoots have emerged

Use Claforan and Vancomycin during shoot induction and after **only** if necessary. Check for contamination daily.
Aim for following conditions:
+23 C
70-90 µmol light, 18/6 light period
Less light and colder temperature might prevent shooting.

| **Root induction plates** | **500 ml** |
|---|---|
| • WPM 2,46 g/L | 1,23 g |
| • MES 0,5 g/L | 0,25 g |
| • sucrose 20 g/L | 10 g |
| • Agar 7 g/L | 3,5 g |
| • set pH 5.6 | |
| • Autoclave | |

**Add when making plates**

- IBA 0,2 mg/L — 100 ul from 1 mg/ml stock

### STOCKS

https://www.sigmaaldrich.com/technical-documents/protocols/biology/growth-regulators.html

| **Auxins** | **Work C** | **Stock C** | **Dissolve** | **Store liquid at** |
|---|---|---|---|---|
| IBA | 0,2 mg/L | 1 mg/ml | 1M NaOH, $dH_20$ | -20°C |
| NAA | 0,2 mg/L | 1 mg/ml | 1M NaOH, $dH_20$ | -20°C |
| **Cytokinins** | | | | |
| BAP | 1 mg/L | 1 mg/ml | 1M NaOH, $dH_20$ | -20°C |
| TDZ | 0,5 mg/L | 1 mg/ml | 1M NaOH, $dH_20$ | -20°C |
| **Antibiotics** | | | | |
| Spectinomycin (sel) | 100 mg/L | 100 mg/ml | | -20°C |
| Rifampicin (broad spec) | 100 mg/L | 50 mg/ml | | -20°C |
| Gentamycin (helper) | 25 mg/L | 10 mg/ml | | +2-8°C |
| Timentin (Agro) | 200 mg/L | 100 mg/ml | | -20°C |
| Vancomycin (Gram+) | 100 mg/L | 100 mg/ml | | -20°C |
| Kanamycin (Plant sel) | 50 mg/L | 100 mg/ml | | -20°C |
| Claforan (Gram-) | 200 mg/L | 250 mg/ml | | -20°C |
| **Other stocks** | | | | |
| Acetosyringone | 20 mg/L | 5 mg/ml | 70% EtOH | +4°C |
| Ascorbic acid | ~750 mg/L | 100 mg/ml | dH20 | -20°C |

**Chemicals**

**PVP:** prevents accumulation of phenolic compounds to toxic concentrations
**Claf:** treat plant tissue infections with Gram-negative bacteria
**Van:** treat plant tissue infections with Gram-positive bacteria.
**Tic:** Timentin is a mixture of ticarcillin and clavµlanic acid. Ticarcillin is a broad spectrum semi-synthetic penicillin. Timentin is used most commonly in the regeneration medium for elimination of the Agrobacterium post-transformation.
**Acetosyringone:** induce the Agro vir-Genes and enhance transformation efficiency

MATERIALS CHECK LIST FOR TRANSFORMATION

- Agro re-suspension media

- Ascorbic acid media

- Co-culture media with PVP

- TDZ, NAA, Ascorbic acid ja Acetosyringone in eppendorfs

- 10ul, 100ul, 1000ul ja 5 ml pipets and tips

- 2 x 50 ml Falcon tubes

- Sterile square plates

- Small syringe needles

- Permanent marker

- Gloves

- Lighter

- Scalpel, tweezers, rack

- Waste containers

- 100% EtOH and container for dipping instruments

- 70% EtOH spray bottle

- Explants ☺

**Prepare these before starting**

| **Agro re-suspension media with PVP** | **100 ml** |
|---|---|
| • MS 2,14 g/L | 0,214 g |
| • Sucrose 20 g/L | 2 g |
| • MES 0,5 g/L | 0,05 g |
| • PVP 10 g/L | 1 g |
| • set pH 5.6 | |
| • Autoclave | |

| **Ascorbic acid media** | **250 ml** |
|---|---|
| • MS 2,14 g/L | 0,535 g |
| • Autoclave | |

**Add before use**

| | |
|---|---|
| • Ascorbic acid ~750 mg/L | 2 ml from 100 mg/ml stock (filter sterilized) |

| **Co-culture media with PVP** | **100 ml** |
|---|---|
| • MS 2,14 g/L | 0,214 g |
| • Sucrose 20 g/L | 2 g |
| • MES 0,5 g/L | 0,05 g |
| • PVP 10 g/L | 1 g |
| • set pH 5.6 | |
| • Autoclave | |

| **Washing media** | **300 ml** |
|---|---|
| • MS 2,14 g/L | 0,65 g |
| • MES 0,5 g/L | 0,15 g |
| • Set pH to 5.6 | |
| • Autoclave | |

| **Selection and shooting media (WPM)** | **500 ml (usually 6x500ml)** |
|---|---|
| • WPM 2,46 g/L | 1,23 g |
| • MES 0,5 g/L | 0,25 g |
| • sucrose 20 g/L | 10 g |
| • Agar 7 g/L | 3,5 g |
| • set pH 5.6 | |
| • Autoclave | |

-Autoclave box of 5 ml tips (step 4).
-Autoclave small sheets of filter paper inside foil (step 6).
-Prepare all chemicals

| | |
|---|---|
| • Ascorbic acid | 100 mg/ml stock (filter sterilized) |
| • Acetosyringone | 5mg/ml stock |
| • Claforan | 250 mg/ml stock |
| • Timentin | 100 mg/ml stock |
| • Vancomycin | 100 mg/ml stock |
| • Kanamycin | 100 mg/ml stock |
| • Rifampicin | 50 mg/ml stock |
| • Gentamycin | 10 mg/ml stock |
| • NAA | 1mg/ml stock |
| • TDZ | 1mg/ml stock |
| • BAP | 1mg/ml stock |
| • IBA | 1 mg/ml stock |
| • LB | |
