## Supplementary material for "*BpLAZY1A* Mediates Transcriptional Polarity and Drives Adaxial Tension Wood–Like Tissue Formation in Silver Birch (*Betula pendula*) Branches": Materials and Methods

**Materials and Methods for**

**Birch genetic material**

The weeping silver birch cultivar *Betula pendula* 'Youngii' carries a premature stop codon in the *BpLAZY1A* gene (Salojärvi et al., 2017). To generate a segregating population, *B. pendula* 'Youngii' (Fig. 1A) was crossed with wild-type pollen (E1970; from the LUKE collection), producing an F1 generation consisting only of wild-type (WT) individuals. A WT tree from the F1 population (Fig. 1B) was subsequently selected and backcrossed with *B. pendula* 'Youngii' to produce a BC1 population ('Youngii' × F1). The BC1 progeny segregated in a 1:1 ratio of WT and mutant phenotypes. The mutant genotype was confirmed by *DpnII* restriction enzyme digestion, which verified the presence of the homozygous SNP associated with the mutant phenotype (Table S1; Fig. S2). Several BC1 lines were propagated in vitro. Over time only lines WT #147, WT #148, and M #161 remained viable under *in vitro* conditions. Of these, WT #147 and M #161 were selected for phenotypic analysis using time-lapse photography.

The silver birch lab clone V5834 (WT) was genetically transformed with the following constructs: *pBpLAZY1A::erVen* and *BpLAZY1A::RNAi*. The *BpLAZY1A* promoter, as well as the *BpLAZY1A* genomic and coding sequences (Table S2), were obtained from the CoGe platform (https://genomevolution.org/coge/).

A 3,633 bp genomic sequence upstream of the *BpLAZY1A* transcription start site (Table S2) was amplified and cloned into the Gateway donor vector P4P1RpGEMt. The donor vectors *pBpLAZY1A* in P4P1RpGEMt, *p221z-erVenusYFP* and *2R3e-nosT2* were subsequently assembled into the destination vector pCAM-kan-R4R3 to generate the *pBpLAZY1A::erVen* construct. Successful transformation of two independent lines was verified using confocal microscopy (Fig. 1 K-N; Fig. S6-6). The *BpLAZY1A::RNAi* construct was generated by amplifying a 210 bp antisense fragment targeting exon 4 (Table S2) from a silver birch cDNA library from the WT (V5834) lab clone. This fragment was cloned into the pDONR-221z donor vector, followed by integration into the destination vector *pK7GWIWG2*, which contains a 35S promoter, 35T terminator, and kanamycin resistance for plant selection. V5834 plants were transformed with this construct, and RNAi functionality was validated through RT-qPCR (Fig 1H). For RT-qPCR analysis, birch RNA was extracted (Table S3) from *in vitro*-grown plants (V5834, *BpLAZY1 RNAi* line 1, and *BpLAZY1 RNAi* line 4). The DNase treated RNA was used for cDNA library synthesis with oligo-dT primers (First Strand cDNA Synthesis Kit, Thermo Scientific) according to the manufacturer´s protocol.

**Silver birch transformation**

All silver birch transformations were conducted with agrobacterium mediated gene transfer according to our lab protocol (SI 7) adapted from Keinonen-Mettälä *et al*. (1998).

**RT-qPCR**

For *BpLAZY1A* RNAi validation, whole in vitro-grown plants without leaves were collected and snap-frozen in liquid $N_2$. For validation of the silver birch bulk RNA-seq data, the same RNA samples were used. Total RNA was extracted using the cetyltrimethylammonium bromide (CTAB) method, followed by DNase treatment and cDNA synthesis as described in Table S3. qRT-PCR analyses were performed in 11 µl reaction volumes using HOT FIREpol® EvaGreen® qPCR Mix Plus (no ROX) 5× reaction mix (Solis BioDyne) on a CFX Opus 384 Real-Time PCR System (Bio-Rad). The PCR program consisted of an initial denaturation step at 95 °C for 12 min, followed by 40 cycles of 95 °C for 15 s, 55 °C for 20 s, and 72 °C for 20 s, with a subsequent melting curve analysis. Three biological replicates were analyzed for each experiment, with three technical replicates included for each biological replicate. Expression levels were normalized against two reference genes, *GapDH1* and *PP2A*. Relative expression levels were calculated using the 2^−ΔΔCt method (Livak and Schmittgen, 2001). All qRT-PCR primers are listed in Table S4. A 1:16 dilution of cDNA was used for bulk RNA-seq validation, whereas a 1:2 dilution was used for *BpLAZY1A* RNAi confirmation

**Phylogenetic analysis of the *LAZY* gene families**

Protein sequences of *Arabidopsis LAZY1-6* were acquired from TAIR (https://www.arabidopsis.org/) and used as a query to identify orthologs in silver birch genome using the blast algorithm in *Betula pendula* genome v1.2 on CoGe (https://genomevolution.org/coge/). LAZY gene sequences previously reported to cause strong shoot phenotypes in rice, maize, *Arabidopsis*, and apple were retrieved from published studies (Li et al., 2007; Dong et al., 2013; Yoshihara et al., 2013; Taniguchi et al., 2017; Dougherty et al., 2023). These sequences were subsequently used as queries in BLAST searches with default settings in Phytozome 14 (https://phytozome-next.jgi.doe.gov/) to identify orthologous LAZY genes from *Oryza sativa* v7.0, *Malus domestica* v1.1, and *Zea mays* RefGen_V4. Protein sequences were aligned using ClustalW with default parameters in MEGA11 (https://www.megasoftware.net/). Redundant sequences were removed prior to a second alignment. Following re-alignment, one *Malus domestica* sequence (MD16G1122900) was excluded, and one *Oryza sativa* sequence (LOC_Os03g29270.1) was trimmed due to the presence of an N-terminal region absent from the other LAZY protein sequences. A phylogenetic tree was constructed using the UPGMA method, and bootstrap support values were calculated from 1,000 replicates.

**Birch time-lapse photography and phenotyping**

Time-lapse photography was performed in a dark room maintained at 20 °C. Illumination was provided by two BX120c7 white LED panels (Valoya) positioned above the plants. Light intensity was approximately 250 µmol $m^{-2}$ $s^{-1}$ at the upper canopy level and 100 µmol $m^{-2}$ $s^{-1}$ at table level. All images were corrected for exposure, contrast, and color balance using Lightroom Classic (Adobe). Videos were compiled from individual image frames using the timeline tool in Photoshop (Adobe), with the frame rate adjusted such that 1 s of video corresponded to 1 d of real time. For the segregating population experiments, plants were grown under an 18 h light / 6 h dark photoperiod. Images captured during the dark period were removed in Lightroom Classic (Adobe) prior to video compilation. Due to inconsistencies arising from abrupt apical movements in the segregating population (linked to frame loss during the dark period) these samples were excluded from the apical movement analysis. V5834 and *BpLAZY1* RNAi plants were grown under continuous light conditions. In all experiments, plants were 3–4 weeks old after potting at the start of filming, and all compared plants were age-matched. Orthogonal movement of branch apices was quantified in ImageJ version 1.54r by measuring deviation of branch tips from the central branch axis over time. Branch curvature in basal internodes was analysed using the Kappa plugin in ImageJ.

**Preparation, staining and imaging of tension wood**

Resin embedded sections (Fig. 3) were prepared by fixation in 4% paraformaldehyde (PFA). Samples were then dehydrated through an ethanol series consisting of 10%, 30%, 50%, 70%, 95%, 100%, and 100% ethanol for 30 min at each step. Samples were subsequently incubated overnight at +4 °C in darkness in a 1:1 mixture of 100% ethanol and Leica Historesin without hardener followed by overnight incubation in Leica Historesin without hardener at +4 °C. Samples were then embedded in a 15:1 mixture of historesin and hardener in plastic molds and polymerized overnight at +20 °C under airtight conditions. Resin blocks were sectioned into 5 µm sections using a microtome. Sections were stained for 30 s with 2% Astra Blue prepared in $dH_2O$ containing 2% glacial acetic acid, followed by thorough rinsing with $dH_2O$. Sections were imaged using a Leica DM6000 microscope with tile-scan acquisition. Individual tile images from each section were subsequently stitched together using the Leica image acquisition software.

Cryo-section samples (Fig S7A-B) were prepared by sampling the basal internode four weeks after decapitation. Adaxial side was first marked and samples put into eppendorfs on ice. Samples were then stored at -20°C. 40 µm thick sections were cut in a cryotome (set to -20°C). Sections were stained on microscope slides with 1% Alcian blue in $dH_2O$ for 10 seconds, washed with $dH_2O$, and subsequently stained with 0,05% Safranin O in 50% EtOH for 10 seconds and washed again with $dH_2O$. Sections were imaged using a Leica DM6000 microscope with tile-scan acquisition. Individual tile images from each section were subsequently stitched together using the Leica image acquisition software.

***In vitro* growth conditions for silver birch**

Silver birch in vitro cultures were maintained under long-day conditions (18 h light / 6 h dark) at approximately 23 °C in Magenta Plant Culture Box containers (PlantMedia). Light intensity was maintained at 50-100 µmol $m^{-2}$ $s^{-1}$. The culture medium consisted of 0.7 g/L MES (Duchefa), 20 g/L sucrose, 7 g/L plant agar (Duchefa), and 3.3 g/L MS medium with vitamins (Duchefa), adjusted to pH 5.7. For root induction, the medium was supplemented with 0.2 mg/L indole-3-butyric acid (IBA; Duchefa), whereas shoot induction medium contained 1 mg/L 6-benzylaminopurine (BAP; Duchefa).

**Silver birch RNA extraction, DNase treatment and cDNA library preparation**

Table S3

**Silver birch gDNA extraction**

Table S5

**Greenhouse growth conditions**

Silver birch trees were cultivated in 3 L pots within greenhouse compartments in the greenhouse installations at the University of Helsinki, Finland. To counteract seasonal variations in outdoor climate, environmental conditions were regulated to maintain a long-day photoperiod (18:6 h light:dark) using Philips MASTER SON-T PIA Plus 400W lamps. Target temperatures were set to 20°C during the day and 18°C at night, while relative humidity at ambient levels (40-60%). The growth substrate consisted of a peat, sand, and vermiculite mixture (6:2:1), fertilized with granular Osmocote Exact (Everris) at a concentration of 2 g/L.

**Silver birch sample preparation and confocal microscopy**

Wild-type (V5834) control and transgenic *pBpLAZY1A::erVen* plants were sampled 1.5–2 months after potting, at a height of 100–130 cm. For branches approximately 30 cm in length, samples were harvested from three positions: the third internode from the apex, and the first and third basal internodes. Main stem samples were isolated from the third apical internode, the mid-stem, and 20 cm above the soil level. All collected tissues were immediately placed in phosphate-buffered saline (PBS) on ice and maintained at 0°C throughout subsequent processing.
Samples were embedded in 4% agarose and cut into 200 µm thick sections using a vibratome. Sections were stained with SR2200 (Renaissance Chemicals) at a 1:1000 dilution in PBS and subsequently imaged using a confocal microscope (Stellaris 8, Leica Microsystems). Excitation wavelengths were set to 405 nm for SR2200 and 515 nm for erVen. Emission was recorded between 427-432 nm for SR2200 channel and between 520-544 nm for erVen channel. Data were acquired either as single optical planes or as z-stacks.

**Bulk RNA-seq**

Plants were sampled two months after potting. Individuals from V5834 and *BpLAZY1A* RNAi lines were selected based on a height profile of 120–160 cm and a main-stem node count of 29–35 internodes. For each plant two branches approximately 30 cm in length displaying similar horizontal growth phenotypes were chosen. The adaxial (upper) side of each branch was marked at three specific positions: the apical segment (10 cm long), the third basal internode (branch mid), and the first basal internode (branch base). Collected samples were immediately snap-frozen in liquid nitrogen and subsequently dissected into adaxial and abaxial fractions within a cryo-chamber maintained at -30°C. Total RNA extraction and DNase treatment were performed as described in Table S3 prior to shipping to Novogene. Five biological replicates of WT and *BpLAZY1A RNAi* samples were selected for mRNA library preparation using poly(A) enrichment. Sequencing was conducted on an Illumina NovaSeq X Plus platform in a paired-end 150 bp (PE150) configuration, yielding approximately 9 G of raw data per sample.

RNA sequencing reads were first processed to remove ribosomal RNA (rRNA) sequences using SortMeRNA v4.3.6 (Kopylova et al., 2012). Quality filtering was performed with Trimmomatic (Bolger et al., 2014), applying the following parameters: LEADING:20, TRAILING:20, SLIDINGWINDOW:3:20, and MINLEN:55. The filtered reads were then aligned to the Betula pendula reference genome (Salojärvi et al., 2017) using STAR aligner v2.7.11b (Dobin et al., 2013) with default settings. Gene-level read counts were generated using featureCounts (Liao et al., 2014) with the parameters "--countReadPairs -t exon" and the publicly available B. pendula gene annotation file (Salojärvi et al., 2017). Differential gene expression analysis was performed using DESeq2 v1.46.0 (Love et al., 2014). The analysis pipeline included normalization, followed by dispersion estimation and statistical testing for differential expression. Pairwise comparisons were conducted using the Wald test. Genes were classified as differentially expressed based on a false discovery rate (FDR) threshold of $p <= 0.1$, fold change ±1 and with p-values adjusted for multiple testing using the Benjamini-Hochberg procedure (Benjamini & Hochberg, 1995). To assess biological relevance, including replicate similarity, we applied principal component analysis (PCA) alongside other visualizations (e.g., heatmaps) produced using custom R scripts. Cell wall and auxin related differentially expressed genes were manually categorized based on the function of their best amino acid sequence blast hits on Araport11 (https://www.arabidopsis.org/tools/blast/).

**Monosaccharide analysis by Trimethylsilyl (TMS) derivatisation**

We analyzed hand ground sample material that was also used for the bulk RNA-seq along the branch axis experiment. Before analyzing the samples, the powdered samples were vacuum freeze dried over a weekend (Savant ModulyoD, Thermo).

**TMS derivatisation**

500 ug (± 5%) dry fine wood powder and 30 ug of inositol, used as internal standard, together with standards of nine monosaccharides (Ara, Rha, Fuc, Xyl, Man, Gal, Glc, GalA and GlcA, each at 5, 10, 20, 50 and 100 ug) were methanolysed and derivatised, and its silylated monosaccharides were separated in GC/MS (7890A/5975C; Agilent Technologies, Santa Clara, CA, USA) according to Sweeley *et al*. (1963 and 1966) and Latha *et al*. (2015).

**Analysis of GC/MS data**

Raw data MS files from GC/MS analysis were converted to NetCDF format in Agilent Chemstation Data Analysis (Version E.02.00.493) and exported to the RDA (version 2016.09; Swedish Metabolomics Centre, Umeå, Sweden). Data pretreatment procedures, such as baseline correction and chromatogram alignment, peak deconvolution and peak integration followed by peak identification was performed in RDA. 4-*O*-methylglucuronic acid was identified according to Chong *et al.* (2013). The amount of each sugar was normalized based on inositol and calculated according to the standards.

**Table S1** Genotypic confirmation of the *Bplazy1a* mutation in the segregating population with RE digestion

gDNA LAZY1 2919 bp residues segregating population genotyping with DpnII

ATGAAGGTAAATTCCTCTGTTCTCCGTTACAAACTTTCTCGATATTAATTCTGCTTGTTAATTTTCATCTACTCTTCTTGCAGTTACTAGGTTGGATGCATCGTAAGTTCCGCCAAAATAGC
TGCGACCCCCTCAAGGATTTTGTCACTGGTAAGAGTACAGGTGCCACCCACATCAAACAAGTTCCTTTAATTTACATTTTATTTTTTATTTTTTTCTTTTGGGTATTAATTCTTGTTCTTG
TGTTTCAGGGCAGCCATCGGTTGAC**GATC**AACAGTACCAGCTAAAGCAAAACTATGGCACCA**GATC**GTTCAAACAAGCCCCAAAAGACCACCACCTTCGGAAGTCTTTCACTGGTC
TAGAAGCAGCAGCAATAGTAGAAGAAGATATTCAAGACTACGGAGAAGAAACATCTGCTGCAATATCTGAGGCGCTCTTCCACGGCTTCCTTGCAATTGGTACTCTGGGTGCAGCA
GAGCAAGTAGTCATCAATGATATTCCGTCAACACCGACATTCGCCATCTCTGTTGAGAATATAACCGAGAAAGAAACGGAAGTGACGGAGAACGATCTGAAGCTGATCAATGATGAG
CTGGAAAAGGTTCTGGGAGCTGATGTACTGAAAGACGATGGTTGCAATTTTTCATCTGGGAGAAACAGCCATGTCAGCACTGGACGAAGTAGTCATGGTAGCACAATTACACTTGG
CGGTAAGCCCATAGAAGGCCCAGATCAGACAAATGGGAATGGAAATGCAGTCTGCCCACTCCAAGGATATCTTTTTGGGTCAGCAATTGAAATGTCAGAAACAACCCCGGTGGCAA
AGAAGGAACATAGAACATCTCTTGGTGAGCTTTTTCAGATGACCAAAATGGCAGAGGAGAATTATTCCGGAACCAAGTGCGATAGGGATCAGGAGAGGCGATCGGAGAAGGAAG
CAGATAAGTCTGCCATGCACTTGATGAAAAAGTTGCTCAAGAAAAAAATGCTCCACGCTTCTTCTCGAACCTCTACAGAGGCTGCTGCAGGTGGAACTCTTGATTCTGCTTCAGCAG
AGAAAAAGCTGCATAAGGTAACTCCTGACTTTGTTATTGTGTTTTTGGTTGATAAATTCCTTATCCTATAATGTGGTGTGGGCTGTAGATACTGCAAATGTTCCACAGGAAAGTTCACC
CTGAAAACTCAACAGCAACTCAAAAGTCTAACAAGACCTACAAGAGTGAAAACAAGAAGAAAATAATGAACAACAATGGGGAATATTACAACAATGGAGATCAGGTGGTGATCCCA
GATGAAGAAATAATTTCATATCCTAAACGAACCCTTTTGAAGGAGAGCATAAGACGCTACAAGAGCCAATCTAACCCGCCACAGTTCACACTTGGCAGCGGTGATTCGAATGGGAAC
AGGGAGCACTGGATCAAAACAGATGCCGAATGTAAGTACACAATATATATACATCTGATCCTCCTCCAAACTCCAAACTCCATTGGCATTGGGAACGCCAACTTTGTGTGTTATCATAA
AAAGAACCACCCGATTGATGATTACAGCACTGTCTTCAATTCTATGCCGGACTTTGGGGCTCTTAATTTTTACAATCGAATATTCCATTACAAATGAAATGAAGAGGAGGGGTTAATTA
AAATAAGAGGGTTAAAATTGTCATTTTATTTTTTTTTGCCCGTCTTGAATAACTAATCTTCCTAATTTGATGCTAATTAATGTAGCATTAATTCCATGCTTTTAAGATTTCGGGCCCGTTTC
TAAACAAAATCTTGAATAGCATTTGTTTGAAAGTGCGATAAAAAAATGCGTTTTTAAAAAATTACTATTTTTTAATATTATTTATTTTGAAATTACAAATCTAAATAAATCCTAATAATTTTTAA
AAATACAGTCAATTATTTGATAAAATTGTAATTTAACATTTAAAATTATGTATTTTTAAGAACATACCCCCTTGTGATTTGAATACGCATATTTTTTTACAAACAAATATCTTCCTAATGGGG
TGCTAATTAATGTAGCATTCGATGCCTTTAAGATTTCACATGCGTTTTTAAACAAAATCCCAAAAAAATGTTTAGTTTGAAAATGCATTTGAACAAAAAAAAAAAAAAAAGGCAGTTTCA
ACATGCAAAAGGCTGAATTGTAATTTTTTAAGTATTTAATTGTGATTTTAAAAATCGTGTTTTCATTAACTACATTTTAAATAAAATGGCTATTTTTTTCAATTTGTGACCTTATGCACATT
CAATACATGGAAAGTAGTGGTGTGACTCAATTAGGATAGCAATGTGATTGCAAAGAGTAATGTTACACACACTTTTACTTTTTCACACTCATTTCTCCATGCATACTCGTAGGTGGCTCT
TAATGTCATTTATGATTAAACAACGTAAACGAAGAATAATTGTGTCATTTAATGTAAATGGCGTAAAAACAAACGACACTAATTAACAACCATGTTTTTTGTAGTGTCATTATAAATCTAA
TAGTAATTTTAAAAGTCACCTAAAAATGAGAGGAAATGGGCATGAGAAAGTGTGAATTCCAATGATTTCAGTACGAAATATCAAAACAAGCCTTGTAAGATGCTTGTTCTGGCGCTTA
TTTGTGATCTAGTAGCTAGCCATCCTGATTGAGTTGTTTGTTTGCATTGTAGATCTTGTGCTGGAGCTCTGAAAGGCAGAAGAAGCAATATTTGAGCGGGGTTCGTCACCCCTGAGGC
TTGAGCAGTTACCAGTGATATCGTCTTCCTTGTTGGTGATGGTTGCTAATAGCATACAATATGTGATAATGAGCTTGGGTTGAACAAATAAATAAAAAGAGAGGAACACCTCTTGTGTC
CGTGAGTGGTCAATGTTGAATGTAAGGTACATGTGTTTTAGTAAGTAATTATATTTCCGCCATTATGGCA

Forward primer(23bp)(BpLAZY1_genot_FWD):TTCTTGTTCTTGTGTTTCAGGGC
Reserve primer(20bp)(BpLAZY1_genot_R30): AGATGGCGAATGTCGGTGTT
Product length: 293bp

**WT (Aa) DpnII digest**:

For A

| | | |
|---|---|---|
| 1(LeftEnd)-DpnII | 1-37 | 37 bp |
| 2 DpnII-DpnII | 38-74 | 37 bp |
| 3 DpnII-(RightEnd) | 75-293 | 219 bp |
| | **3 fragments: 37 bp, 37 & 219 bp** | |

For a

| | | |
|---|---|---|
| 1(LeftEnd)-DpnII | 1-37 | 37 bp |
| 4 DpnII-(RightEnd) | 38-293 | 256 bp |

| | |
|---|---|
| For Aa | **3 fragments: 37 bp & 219 bp & 256 bp** |

**Homozygote mutant DpnII digest**:

| # Ends | Coordinates | Length (bp) |
|---|---|---|
| 1 (LeftEnd)-DpnII | 1-37 | 37 bp |
| 4 DpnII-(RightEnd) | 38-293 | 256 bp |
| | **2 fragments: 37 bp & 256 bp** | |

**Table S2** ***BpLAZY1A*** **promoter, gDNA and cDNA sequences**

*BpLAZY1A* promoter sequence 3633 bp (Bpev01.c0052.g0076.m0001). Primer sites are highlighted with blue.

GTTAAAGAGAGTTAAAGACATTGGGACAAACAGCCCCAATGCCCCCATAACATTATGTCCAAAAAAACAAGACGTAGTCATATCATATCACATCATATAGTGTGGGGAATGGGAATCGATCCAGTG
CTCCTACATCTAATTCTGCACTATCACATCTTGGTGAGGTGTTCCCTGCAGAGTCGGCTCCCCTCATGTTAGCGGTAGAAACTATCACATTACTTCGTTTTTTAGTTTCATGTTTGCTTCCAAAAAGAG
GACCCCCACACGTTGAGTTTATCGCCATTCTTTTCATTCAATCTAGCCTTTTTTTTTTTCTTTTTTTCTTTTATGATCAAATGTCAGTGTAGCATCATTTACATATCTTTTTGCTTTCCTGACATAACTGAA
TATTGTTGTCTGACATATCGGTCATACAAGTGTGTGTGTCTTTTTTTTTTTCCTTTAAAAAGTAATGTTATCATTCACACCACAATTGCTAACGTGTTAAAGGCTAATAGTCATTGATTAATATTTTTATTT
TCTTTTTTAACAAGGGCTAATTGAAGGACTAATAGCCTATAACCACCACGCCAGCATAGTGTGTGTTGATCATTAGGTTAGATGTATAGTGTAATTTTTTTTTTTTTTTTTTAACAATGACTAAACCTAT
TTTTATCAGGGTTAATTAGGTGGGATGTATTTAGTTATGGGTATTATGTTCTTTAATTATTAGATCTCTCCTCCTACTAGTACTAAACTCTAAATTAAATAACTATATATATATGTTACGGGTGGGCTCAAT
TTTTGTCTCAAATATAAGTCGATAGAAAATAATAAATTTAATTATTTAATTAATACTTTCATAACACAATTTATTTTTCATTAGTTTCGGCTAGATCATCGGACTTGTTTGTGTTTGTTTCAATGTCGGTAG
ATAATATGTTATTCTTGGGATACTAGGACAATATATATTATCAAAACTTGCATGATTCGACAGGTTACACAGGCCATCCATACTCATCTTTTCAACATGATGCAATGTTTCGTGGTATGTGGTCCAATTTT
AAATATCTGCCTTGTGCTATTTCATTTACTAAATTATTATCAACCAGGAAAGTAAACTATTGCTCAATCATAAAGTTGAAATCTATCATAGTGATGGAAAAAAGAGATAAAAGAATGGGAGCAGGCCA
ATCTATCAGTAGCTAGTACAATTATTAGGGTTGAAATCGGTACTTTTTTAAGCATTATCTGTTTATCTCAAGCTCTTAAATTCTCATGCCATAGAATATATAGTTGAAAAAAAAACAAAGGAACAGAAT
CACTGAAATAGAAAGATCATGACAATTATTAAGCAGCTTCTAGTTAACATTCTCGATCTCTTTTCTCATTCTCTATCAACTTCCTCTCAAAAATAGATAAATTAAGAAGCATAATTTGCTTGATACCTCT
GGAGATCATCCTATCTCACTCATTGGCCAAATTAAGGAACTTTTAACAGGACCAGCTGGTTTACATCCATTGTTTGTTATCCTGAGTGTGTGTGTTTATGTGTTTGTTTGTAATTGAACTTGCTCATGG
GACCAGGTTTACATCCATTGTTCATTGAAACGTGGGTTATAGCTTGAGTGGTCTTTTCTATATAATTACTGGCTAGAATCGCGAGGAGTAAGTTTCTCCTCGTCAAGGATAATGCCTGCGTCACCTTT
TGTCTCTAAGCCTGCAACCCTTTTCTCTTTTGCTGTGCACTTTCTCTTCTTGAAGATGCTCGATATATCGTGGGACACGCTTAACTTCTCATCAAGCCCAACAATCCTGCTCCAGGACCAGATGTACTC
TTCACAGAAAGAACACACTTCTGACCATAATCATTAAGAAACACTGCTTCATCACATCTACCAACTTTAATTACATTCCACCGCAGTCACAAAGCTTTGTAATTACCAATTGCAACTAATTCCTTACAA
TGTCAGTACTATGAGGCACAAGTTTAGGGTAGTTGGTAATTTGTATATTTATGTGTCGGATTAGGGTTGTCGAGGTATAGGTATTTATAAAATTATATAGGCCAACACTAATTTGATTCATTGAATTAAA
CGGGTCACACTTTTTAATTCTAATTCATTAATTTCATATTTAATTCATATCAAATTCGCAAGTTGTGTCGAGACATGAGTATAACACTTATAAGTTAATCTTAACTTGACTTATTTAGTTAAACAAATTAA
ACCTGTTAACTGTAACTAATTAATTATTTTGTGTCGAATTAGTGAATCATGTCAAAAATTGTGAGTTCAAAGTAGAACACATAACAATTGCTGAAAAAATAATATACAAGTGCAGTAGATTGGGACTA
AAAGAGCTGTTTTAATTCGCCAACCATCGATACCTGCACGTTATACCAGATCGAATAAGTACTTGGCCGGCTGAAACAAAAGAATATGTACCAGATATTCCAGGTAGTACTTCATGGGCATCCAATTA
TTATTATTATTATTATTGTAAGTATGGGCATCCAATTATTGCTTCTGCAGGATTAAGATGGTTGCGCACCGGCCGAAACATAATAATTCAAACAATTATATATTTAGATTTGACTAAACATAACCCCGACT
TATATATTATTGTTAATCAAATGGGTTCCTTCAGTTAGTCAGAAGTACATCGTTGGACCTACTGGAAAAAGCATAGGGCAATCAAACTCGAACTTTTGTGTCGACCCAGATGAGAAAAGATCTACTTC
ATTGTCATACATATATATCGATGATCGGCCTGACAAAATATTCTCTAAGAGTCCCCACAATTCAGTGTGCATGGATACTCCTACCAAATACCCAATGAAATAATCATCATTACCTCATGCAAAAAGTGCA
TGTGCACGAAAAATGGCAGGTTTTGGTCACATACATCGATCCAAACGTGCTTACTTTTTCTTTTTTTATAGGCCAAGTTGTGAATAAATTAAAAAGAAGATTTGAAAAATACGAGTGATATTCTTCGC
TAAGTATCAGCAAAATGGGGTTATACTGCAGTACATGTGTATTATAAGTTGATGGGCAGTTGGGCACCCTTTCCTTTGCAAGCCAATTTATAAGAAGGTTGCAAATTTTGAGTAGAACTTGCTCTTA
AAAACTGACTTATAATTAAATAGAGAGCGTCTAAAAACTTATATACATATGGAAGACTTGGGATTGTAATTAATTAACATCAGGAAAATGAAAAAGAAAAAGAAAAAGAAAAACTGATATATGCATT
TTTCTGTGCTGAGTTGCATCTTGAAATACGCGTACAGAGAAAGTACTGAGAAATATGACTCATTCAAGAACACCAAATCAAGAAAAGAAGAACAAGAAAAATTGAACTCTACGATGAACAGCAGC
TAATTGCTGAGTGAGAATATAGGATAAGCAGCTAATTGGGGAGGGGGGTCCAGGACCCACTCTGCATTTGGCCAGTTGATTTTATCAGGACAAAGCATTGCCAACATGAATTTGTAATCATCTGTG
TGTTTCTTTCTACCTACTTTCATTAACTGTTTGACATCTTATTCATTAACAAGCACACAGCTATGTATGAAAACATGATCACCCTCTCACATAACTAAACTACTGATCGATCATCAAAAACTGCAAGTTT
CTGAGACACCATTCAAAG

*BpLAZY1A* gDNA sequence 2686 bp (Bpev01.c0052.g0076.m0001). Exons highlighted with green. Fragment used for BpLAZY1A RNAi highlighted with blue. Primers for verifying BpLAZY1A RNAi with qPCR highlighted with red.

ATGAAGGTAAATTCCTCTGTTCTCCGTTACAAACTTTCTCGATATTAATTCTGCTTGTTAATTTTCATCTACTCTTCTTGCAGTTACTAGGTTGGATGCATCGTAAGTTCCGCCAAAATAGCTGCGACCCCCTCAAGGATTTTGTCACTGG
TAAGAGTACAGGTGCCACCCACATCAAACAAGTTCCTTTAATTTACATTTTATTTTTTATTTTTTTCTTTTGGGTATTAATTCTTGTTCTTGTGTTTCAGGGCAGCCATCGGTTGACGATCAACAGTACCAGCTAAAGCAAAACTATGGC
ACCAGATCGTTCAAACAAGCCCCAAAAGACCACCACCTTCGGAAGTCTTTCACTGGTCTAGAAGCAGCAGCAATAGTAGAAGAAGATATTCAAGACTACGGAGAAGAAACATCTGCTGCAATATCTGAGGCGCTCTTCCACGGCTT
CCTTGCAATTGGTACTCTGGGTGCAGCAGAGCAAGTAGTCATCAATGATATTCCGTCAACACCGACATTCGCCATCTCTGTTGAGAATATAACCGAGAAAGAAACGGAAGTGACGGAGAACGATCTGAAGCTGATCAATGATGAGC
TGGAAAAGGTTCTGGGAGCTGATGTACTGAAAGACGATGGTTGCAATTTTTCATCTGGGAGAAACAGCCATGTCAGCACTGGACGAAGTAGTCATGGTAGCACAATTACACTTGGCGGTAAGCCCATAGAAGGCCCAGATCAGAC
AAATGGGAATGGAAATGCAGTCTGCCCACTCCAAGGATATCTTTTTGGGTCAGCAATTGAAATGTCAGAAACAACCCCGGTGGCAAAGAAGGAACATAGAACATCTCTTGGTGAGCTTTTTCAGATGACCAAAATGGCAGAGGAG
AATTATTCCGGAACCAAGTGCGATAGGGATCAGGAGAGGCGATCGGAGAAGGAAGCAGATAAGTCTGCCATGCACTTGATGAAAAAGTTGCTCAAGAAAAAAATGCTCCACGCTTCTTCTCGAACCTCTACAGAGGCTGCTGCAG
GTGGAACTCTTGATTCTGCTTCAGCAGAGAAAAAGCTGCATAAGGTAACTCCTGACTTTGTTATTGTGTTTTTGGTTGATAAATTCCTTATCCTATAATGTGGTGTGGGCTGTAGATACTGCAAATGTTCCACAGGAAAGTTCACCCTG
AAAACTCAACAGCAACTCAAAAGTCTAACAAGACCTACAAGAGTGAAAACAAGAAGAAAATAATGAACAACAATGGGGAATATTACAACAATGGAGATCAGGTGGTGATCCCAGATGAAGAAATAATTTCATATCCTAAACGAACC
CTTTTGAAGGAGAGCATAAGACGCTACAAGAGCCAATCTAACCCGCCACAGTTCACACTTGGCAGCGGTGATTCGAATGGGAACAGGGAGCACTGGATCAAAACAGATGtaCCGAATGTAAGTACACAATATATATACATCTGATCC
TCCTCCAAACTCCAAACTCCATTGGCATTGGGAACGCCAACTTTGTGTGTTATCATAAAAAGAACCACCCGATTGATGATTACAGCACTGTCTTCAATTCTATGCCGGACTTTGGGGCTCTTAATTTTTACAATCGAATATTCCATTACA
AATGAAATGAAGAGGAGGGGTTAATTAAAATAAGAGGGTTAAAATTGTCATTTTATTTTTTTTTGCCCGTCTTGAATAACTAATCTTCCTAATTTGATGCTAATTAATGTAGCATTAATTCCATGCTTTTAAGATTTCGGGCCCGTTTCTA
AACAAAATCTTGAATAGCATTTGTTTGAAAGTGCGATAAAAAATGCGTTTTTAAAAATTACTATTTTTTAATATTATTTATTTTGAAATTACAAATCTAAATAAATCCTAATAATTTTTAAAAATACAGTCAATTATTTGATAAAATTGTAATT
TAACATTTAAAATTATGTATTTTTAAGAACATACCCCCTTGTGATTTGAATACGCATATTTTTTTACAAACAAATATCTTCCTAATGGGGTGCTAATTAATGTAGCATTCGATGCCTTTAAGATTTCACATGCGTTTTTAAACAAAATCCCAA
AAAATGTTTAGTTTGAAAATGCATTTGAACAAAAAAAAAAAAAAAAGGCAGTTTCAACATGCAAAAGGCTGAATTGTAATTTTTTAAGTATTTAATTGTGATTTTAAAAATCGTGTTTTCATTAACTACATTTTAAATAAAATGGCTATTT
TTTTCAATTTGTGACCTTATGCACATTCAATACATGGAAAGTAGTGGTGTGACTCAATTAGGATAGCAATGTGATTGCAAAGAGTAATGTTACACACACTTTTACTTTTTCACACTCATTTCTCCATGCATACTCGTAGGTGGCTCTTAAT
GTCATTTATGATTAAACAACGTAAACGAAGAATAATTGTGTCATTTAATGTAAATGGCGTAAAAACAAACGACACTAATTAACAACCATGTTTTTTGTAGTGTCATTATAAATCTAATAGTAATTTTAAAAGTCACCTAAAAATGAGAGG
AAATGGGCATGAGAAAGTGTGAATTCCAATGATTTCAGTACGAAATATCAAAACAAGCCTTGTAAGATGCTTGTTCTGGCGCTTATTTGTGATCTAGTAGCTAGCCATCCTGATTGAGTTGTTTGTTTGCATTGTAGATCTTGTGCTGG
AGCTCTGA

BpLAZY1A CDS 1209 bp (Bpev01.c0052.g0076.m0001). SNP(C->A) in *B. pendula* ´Youngii´ highlighted with red.

ATGAAGTTACTAGGTTGGATGCATCGTAAGTTCCGCCAAAATAGCTGCGACCCCCTCAAGGATTTTGTCACTGGGCAGCCATCGGTTGACGATCAACAGTACCAGCTAAAGCAAAACTATGGCACC
AGATCGTTCAAACAAGCCCCAAAAGACCACCACCTTCGGAAGTCTTTCACTGGTCTAGAAGCAGCAGCAATAGTAGAAGAAGATATTCAAGACTACGGAGAAGAAACATCTGCTGCAATATCTGA
GGCGCTCTTCCACGGCTTCCTTGCAATTGGTACTCTGGGTGCAGCAGAGCAAGTAGTCATCAATGATATTCCGTCAACACCGACATTCGCCATCTCTGTTGAGAATATAACCGAGAAAGAAACGGA
AGTGACGGAGAACGATCTGAAGCTGATCAATGATGAGCTGGAAAAGGTTCTGGGAGCTGATGTACTGAAAGACGATGGTTGCAATTTTTCATCTGGGAGAAACAGCCATGTCAGCACTGGACG
AAGTAGTCATGGTAGCACAATTACACTTGGCGGTAAGCCCATAGAAGGCCCAGATCAGACAAATGGGAATGGAAATGCAGTCTGCCCACTCCAAGGATATCTTTTTGGGTCAGCAATTGAAATGT
CAGAAACAACCCCGGTGGCAAAGAAGGAACATAGAACATCTCTTGGTGAGCTTTTTCAGATGACCAAAATGGCAGAGGAGAATTATTCCGGAACCAAGTGCGATAGGGATCAGGAGAGGCGAT
CGGAGAAGGAAGCAGATAAGTCTGCCATGCACTTGATGAAAAAGTTGCTCAAGAAAAAAATGCTCCACGCTTCTTCTCGAACCTCTACAGAGGCTGCTGCAGGTGGAACTCTTGATTCTGCTTCA
GCAGAGAAAAAGCTGCATAAGATACTGCAAATGTTCCACAGGAAAGTTCACCCTGAAAACTCAACAGCAACTCAAAAGTCTAACAAGACCTACAAGAGTGAAAACAAGAAGAAAATAATGAAC
AACAATGGGGAATATTACAACAATGGAGATCAGGTGGTGATCCCAGATGAAGAAATAATTTCATATCCTAAACGAACCCTTTTGAAGGAGAGCATAAGACGCTACAAGAGCCAATCTAACCCGCCA
CAGTTCACACTTGGCAGCGGTGATTCGAATGGGAACAGGGAGCACTGGATCAAAACAGATGCCGAATATCTTGTGCTGGAGCTCTGA

**Table S3**

RNA-extraction for birch samples. RIN quality has always been >8 with this method. Method adapted from (Chang et al. 1993).

**Solutions for extraction:**

- **β-mercaptoethanol:** 15 µl per 750 µl extraction buffer
- **EtOH 70 % (-20°)**
- **Chloroform:Isoamylalcohol (24 : 1)**
- **RNase free $H_2O$**
- **Extraction buffer (50 ml)**. Can be prepared in advance. Stays good for months.

| | |
|---|---|
| 2% CTAB | 1g |
| 100mM Tris-Hcl (pH 8.0) | 5 ml (1 M) |
| 25 mM EDTA | 2,5 ml (0,5 M) |
| 2 M NaCl | 20 ml (5 M) |
| 2% PVP40 | 1g |
| RNase free $H_2O$ | fill to 50 ml |

**RNA extraction, DNase treatment, concentration measurement with Qubit**
Note! Keep samples at -80°C.

**Day 1**

1. Grind samples into very fine powder in liquid nitrogen with mortar and pestle. Weight 40-100 mg of sample per 2 ml tube. Keep samples in liquid nitrogen at all possible times. When grinding is finished, keep lid open for 1 min after placing tubes into cold block. Otherwise, they will explode. Grinding can be done before extraction.
2. Set heat block to 65°C in fume hood
3. Put extraction buffer + β-mercaptoethanol (750 µl + 15 µl) into separate 2 ml Eppendorfs, vortex and place in the heat block.
4. Prepare a set of 2 ml sterile-RNA eppendorfs with ~750 µl chloroform:isoamylalcohol (24:1) keep them at room temperature in the fume hood.
5. Just before extraction, take samples from -80°C to -20°C. Take 4 samples at a time and put into a room temperature tube tray for few minutes in the fume hood.
6. Add 750 µl 65°C (extraction buffer + β-mercaptoethanol solution). Vortex thoroughly and make sure no clumps remain. Leave tubes in 65°C for 3 min**.** Vortex again.
7. Add 750 µl chloroform:isoamylalcohol (24:1), mix by inverting 5 seconds.
8. Separate phases in a bench-top centrifuge full speed (13000 rpm) at **RT for 10 min**. Pick **upper layer (500-750 ul)** to the prepared new 2 ml tube containing 750 µl Chloroform:Isoamylalcohol (24:1) and mix by inverting for 5 seconds. Discard lower phase in chloroform:isoamylalcohol waste.
9. Separate phases at full speed (13000 rpm) at **RT for 10 min.** Pick upper layer **(400 µl)** without disturbing phases into a 1,5 ml Eppendorf and add **100 µl** of ice cold 10 M LiCl.
10. Mix and precipitate RNA in -20°C overnight.

**Day 2**

11. Centrifuge (13000 rpm) at **+4°C for 15 min**. Pour out supernatant.
12. Pipette 1 ml of -20°C 70% EtOH (EtOH diluted with RNase free water).
13. Centrifuge again (13000 rpm) at **+4°C for 10 min**. Pour away and pipette out supernatant.
14. Let RNA dry for **10 min** (longer drying time will degrade RNA)
15. Add **25-50 ul of RNase free** $H_2O$. You can add more water if you know that the RNA concentration is very high, less if you have minute starting material.
16. Mix tubes on ice on shaker for 1h. Then mix by hand to dissolve any remaining pellet.
17. Measure concentration with nanodrop and dilute samples to 200 ng/ul ±10%.
18. Proceed with Ambien DNA-*free* kit (or equivalent) according to the protocol.
19. After DNase treatment, measure RNA concentration with Qubit.
20. Finally check RIN with Bioanalyzer, if required for downstream analysis.
21. When applicable, the DNase treated RNA (1 µg) was used for cDNA library synthesis with oligo-dT primers according to the manufacturer´s protocol (First Strand cDNA Synthesis Kit, Thermo Scientific).

**Table S4** Primers used in this study

| primer | Gene ID | Gene name | Primer sequence (5´ to 3´) | Tm (C˚) | Product length (bp) | usage | reference |
|---|---|---|---|---|---|---|---|
| BpLAZY1A genotype F | Bpev01.c0052.g0076 .m0001 | *BpLAZY1A* | GACATCTTATTCATTAACAAGCA C | 58,3 | 1376 | Youngii genotyping | Muranen *et al.,* 2026 |
| BpLAZY1A genotype R | Bpev01.c0052.g0076 .m0001 | *BpLAZY1A* | CAATATGTGATAATGAGCTTGGG T | 62,7 | 1376 | Youngii genotyping | Muranen *et al.,* 2026 |
| AttB4-F2 pBpLAZY1A | Bpev01.c0052.g0076 .m0001 | *BpLAZY1A* | GGGGACAACTTTGTATAGAAAA GTTGTTGTTAAAGAGAGTTAAA GACATTGG | 56,9 | 3633 | cloning of BpLAZY1A promoter | Muranen *et al.,* 2026 |
| AttB1-R1 pBpLAZY1A | Bpev01.c0052.g0076 .m0001 | *BpLAZY1A* | GGGGACTGCTTTTTTGTACAAAC TTGCCTTTGAATGGTGTCTCAGA AAC | 60,2 | 3633 | cloning of BpLAZY1A promoter | Muranen *et al.,* 2026 |
| attB1-F1 BpLAZY1A RNAi | Bpev01.c0052.g0076 .m0001 | *BpLAZY1A* | GGGGACAAGTTTGTACAAAAAA GCAGGCTCTGTTCCCATTCGAAT CAC | 60,5 | 205 | cloning of BpLAZY1A RNAi | Muranen *et al.,* 2026 |
| attB2-R1 BpLAZY1A RNAi | Bpev01.c0052.g0076 .m0001 | *BpLAZY1A* | GGGGACCACTTTGTACAAGAAA GCTGGGTGTCTAACAAGACCTA CAAGAGTG | 56,3 | 205 | cloning of BpLAZY1A RNAi | Muranen *et al.,* 2026 |
| BpLAZY1A F1 qPCR | Bpev01.c0052.g0076 .m0001 | *BpLAZY1A* | GAAGGAGAGCATAAGACGC | 59,2 | 130 | BpLAZY1A RNAi qPCR | Muranen *et al.,* 2026 |
| BpLAZY1A R1 qPCR | Bpev01.c0052.g0076 .m0001 | *BpLAZY1A* | TCAGAGCTCCAGCACAAG | 60,5 | 130 | BpLAZY1A RNAi qPCR | Muranen *et al.,* 2026 |
| GapDH1 F1 | Bpev01.c1040.g0016 .m0001 | *GapDH1* | AGAATACAAGCCAGAACTCAAC | 58,9 | 188 | LAZY1A RNAi qPCR reference | Moschenskaya *et al.*, 2021 |
| GapDH1 R1 | Bpev01.c1040.g0016 .m0001 | *GapDH1* | CTCTACCACCTCTCCAATCC | 60,1 | 188 | LAZY1A RNAi qPCR reference | Moschenskaya *et al.*, 2021 |
| PP2A F1 | Bpev01.c0088.g0042 .m0001 | *PP2A* | GAGGATAGGCATTGGAGAG | 58,8 | 210 | LAZY1A RNAi qPCR reference | Sutela *et al.*, 2011 |
| PP2A R1 | Bpev01.c0088.g0042 .m0001 | *PP2A* | GCATCACGGATCGAGTAA | 59,8 | 210 | LAZY1A RNAi qPCR reference | Sutela *et al.*, 2011 |
| COBL4 F1 | Bpev01.c0189.g0059 .m0001 | *COB3* | AGCATCCCAATCTCAACAAT | 61 | 240 | Bp RNA-seq GOI qPCR | Muranen et al. 2026 |
| COBL4 R1 | Bpev01.c0189.g0059 .m0001 | *COB3* | CGTCGCCATTGAAGTAAACT | 60 | 240 | Bp RNA-seq GOI qPCR | Muranen et al. 2026 |
| CESA8 F1 | Bpev01.c0374.g0018 .m0001 | *CSL* | AGCTTTTCTATGCCTTCTGG | 60 | 233 | Bp RNA-seq GOI qPCR | Muranen et al. 2026 |
| CESA8 R1 | Bpev01.c0374.g0018 .m0001 | *CSL* | TGTATGGTTGGAAAATTCTGG | 61 | 233 | Bp RNA-seq GOI qPCR | Muranen et al. 2026 |
| IAA29 F1 | Bpev01.c0126.g0009 .m0001 | *IAA13* | GTACGTGAAGGTGAAGATGGA | 62 | 227 | Bp RNA-seq GOI qPCR | Muranen et al. 2026 |
| IAA29 R1 | Bpev01.c0126.g0009 .m0001 | *IAA13* | GCACAGACCCAATGAAAGT | 60 | 227 | Bp RNA-seq GOI qPCR | Muranen et al. 2026 |
| SAUR20 F1 | Bpev01.c1577.g0006 .m0001 | *SAUR* | CTATTCGTTTGCCTGGTATTACA | 62 | 271 | Bp RNA-seq GOI qPCR | Muranen et al. 2026 |
| SAUR20 R1 | Bpev01.c1577.g0006 .m0001 | *SAUR* | GAAGTGAGATCAATGAAGTCGT C | 62 | 271 | Bp RNA-seq GOI qPCR | Muranen et al. 2026 |

**Table S5 Silver birch gDNA extraction with EZNA SP Plant DNA kit (modified for birch)**

SP3 and SPW Buffers must be diluted with 100% ethanol before use

1. Weigh 100-120 mg of **young leaf material** and grind with a mortar and pestle in liquid nitrogen (tissue homogenizer can be used also). It is very important to use young leaves. Mature leaves contain secondary metabolites that inhibit PCR reactions.
2. Put ground material into 2 ml tube.
3. Add 650 µl SP1 buffer and 5µl Rnase A.
4. Mix thoroughly (no clumps).
5. Incubate 1-2 hours at 65°C. Mix samples every 30 min during incubation by inverting the tube.
6. Add 210 µl SP2 buffer and mix thoroughly.
7. Keep tubes on ice for 5 minutes.
8. Put DNase free $H_2O$ to 65°C, 100 µl / sample (step 28)
9. Centrifuge samples for 10 minutes at 13 000 rpm
10. Transfer supernatant to the Homogenizer Mini Column (green) with a collection tube.
11. Centrifuge 2 min at 13 000 rpm
12. Transfer cleared lysate to a 2 ml tube and mix with 810 µl of SP3 buffer. Do not disturb or transfer any of the insoluble pellets.
13. Transfer 725 µL of the sample to the HiBind® DNA Mini Column (blue) with a collection tube.
14. Centrifuge 1 min at 13 000 rpm.
15. Discard filtrate and reuse the collection tube.
16. Transfer rest of the sample to the HiBind® DNA Mini Column (blue).
17. Centrifuge 1 min at 13 000 rpm.
18. Discard filtrate and the collection tube.
19. Transfer the HiBind® DNA Mini Column (blue) to a new 2 mL Collection Tube.
20. Add 650 µL SPW Buffer.
21. Centrifuge 1 min at 13 000 rpm.
22. Discard filtrate and reuse the collection tube.
23. Add 650 µL SPW Buffer.
24. Centrifuge 1 min at 13 000 rpm.
25. Discard filtrate and reuse the collection tube.
26. Centrifuge 1 min at 13 000 rpm to remove residual ethanol. Discard collection tube.
27. Transfer the HiBind® DNA Mini Column (blue) into a clean 1.5ml microcentrifuge tube.
28. Add 100 µl of preheated (65°C) DNase free $H_2O$.
29. Incubate for 5 min at room temperature.
30. Centrifuge 1 min at 8 000 rpm at RT.
31. Measure DNA concentration and purity with Nanodrop.
32. Store eluted DNA at -20°C.
